# CalmAn: An open source tool for scalable Calcium Imaging data Analysis

**DOI:** 10.1101/339564

**Authors:** Andrea Giovannucci, Johannes Friedrich, Pat Gunn, Jérémie Kalfon, Sue Ann Koay, Jiannis Taxidis, Farzaneh Najafi, Jeffrey L. Gauthier, Pengcheng Zhou, David W. Tank, Dmitri Chklovskii, Eftychios A. Pnevmatikakis

**Affiliations:** Center for Computational Biology, Flatiron Institute, New York, USA; Department of Statistics and Center for Theoretical Neuroscience, Columbia University, New York, USA; ECE Paris, Paris, France; Princeton Neuroscience Institute, Princeton University, New Jersey, USA; Department of Neurology, UCLA, California, USA; Cold Spring Harbor Laboratory, New York, USA

## Abstract

Advances in fluorescence microscopy enable monitoring larger brain areas *in-vivo* with finer time resolution. The resulting data rates require reproducible analysis pipelines that are reliable, fully automated, and scalable to datasets generated over the course of months. Here we present CaImAn, an open-source library for calcium imaging data analysis. CaImAn provides automatic and scalable methods to address problems common to pre-processing, including motion correction, neural activity identification, and registration across different sessions of data collection. It does this while requiring minimal user intervention, with good performance on computers ranging from laptops to high-performance computing clusters. CaImAn is suitable for two-photon and one-photon imaging, and also enables real-time analysis on streaming data. To benchmark the performance of CaImAn we collected a corpus of ground truth annotations from multiple labelers on nine mouse two-photon datasets. We demonstrate that CaImAn achieves near-human performance in detecting locations of active neurons.

## Introduction

Understanding the function of neural circuits is contingent on the ability to accurately record and modulate the activity of large neural populations. Optical methods based on the fluorescence activity of genetically encoded calcium binding indicators (*Chen et al., 2013*) have become a standard tool for this task, due to their ability to monitor *in vivo* targeted neural populations from many different brain areas over extended periods of time (weeks or months). Advances in microscopy techniques facilitate imaging larger brain areas with finer time resolution, producing an ever-increasing amount of data. A typical resonant scanning two-photon microscope produces data at a rate greater than 50GB/Hour^1^, a number that can be significantly higher (up to more than 1TB/Hour) with other custom recording technologies (*Sofroniew et al*. (*2016*); *Ahrens et al*. (2013); *Flusberg et al. (2008)*; *Cai et al*. (*2016*); *Prevedel et al. (2014); Grosenick et al. (2017); Bouchard et al. (2015))*.

This increasing availability and volume of calcium imaging data calls for automated analysis methods and reproducible pipelines to extract the relevant information from the recorded movies, i.e., the locations of neurons in the imaged Field of View (FOV) and their activity in terms of raw fluorescence and/or neural activity (spikes). The typical steps arising in the processing pipelines are the following (Fig. 1 a): i) Motion correction, where the FOV at each data frame (image or volume) is registered against a template to correct for motion artifacts due to the finite scanning rate and existing brain motion, ii) source extraction where the different active and possibly overlapping sources are extracted and their signals are demixed from each other and from the background neuropil signals (Fig. 1 b), and iii) activity deconvolution, where the neural activity of each identified source is deconvolved from the dynamics of the calcium indicator.

**Figure 1.**
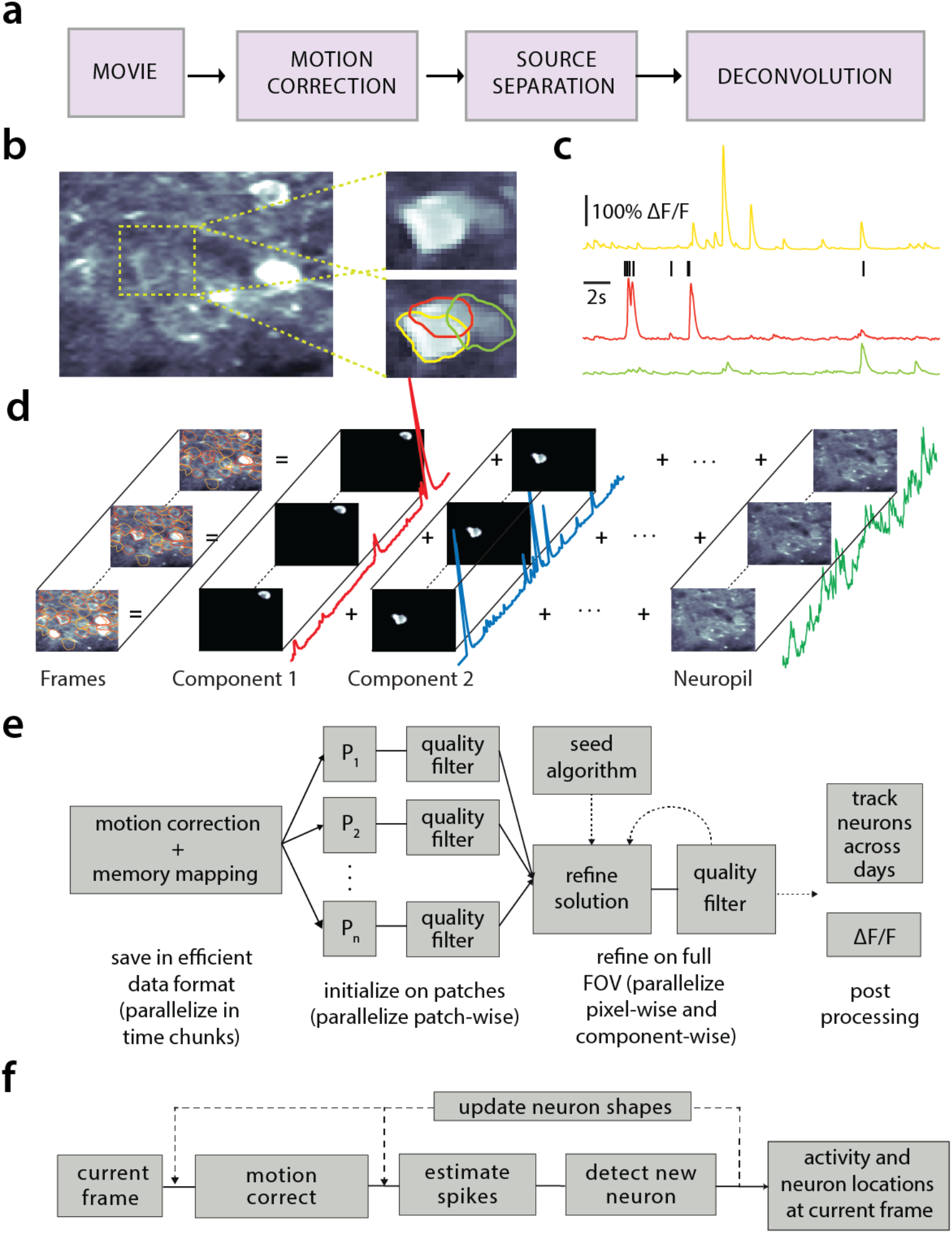
Processing pipeline of CaImAn for calcium imaging data. (a) The typical pre-processing steps include (i) correction for motion artifacts, (ii) extraction of the spatial footprints and fluorescence traces of the imaged components, and (iii) deconvolution of the neural activity from the fluorescence traces. (b) Time average of 2000 frames from a two-photon microscopy dataset (left) and magnified illustration of three overlapping neurons (right), as detected by the CNMF algorithm. (c) Denoised temporal components of the three neurons in (b) as extracted by CNMF and matched by color (in relative fluorescence change, Δ*F/F*). (d) Intuitive depiction of CNMF. The algorithm represents the movie as the sum of rank-one spatio-temporal components capturing either neurons and processes, plus additional non-sparse low-rank terms for the background fluorescence and neuropil activity. (e) Flow-chart of the CaImAn batch processing pipeline. From left to right: Motion correction and generation of a memory efficient data format. Initial estimate of somatic locations in parallel over FOV patches using CNMF. Refinement and merging of extracted components via seeded CNMF. Removal of low quality components. Final domain dependent processing stages. (f) Flow-chart of the CaImAn online algorithm. After a brief mini-batch initialization phase, each frame is processed in a streaming fashion as it becomes available. From left to right: Correction for motion artifacts. Estimate of activity from existing neurons, identification and incorporation of new neurons. Periodically, the spatial footprints of inferred neurons are updated (dashed lines).

### Related work

#### Source extraction

Some source extraction methods attempt the detection of neurons in static images using supervised or unsupervised learning methods. Examples of unsupervised methods on summary images include graph-cut approaches applied to the correlation image (*Kaifosh et al., 2014; Spaen et al., 2017*), and dictionary learning (*Pachitariu et al., 2013*). Supervised learning methods based on deep neural networks have also been applied to the problem of neuron detection (*Apthorpe et al., 2016; Klibisz et al., 2017*). While these methods can be efficient in detecting the locations of neurons, they cannot infer the underlying activity nor do they readily offer ways to deal with the spatial overlap of different components.

To extract temporal traces together with the spatial footprints of the components one can use methods that directly represent the full spatio-temporal data in a matrix factorization setup e.g., independent component analysis (ICA) (*Mukamel et al., 2009*), constrained nonnegative matrix factorization (CNMF) (*Pnevmatikakis et al., 2016*) (and its adaptation to one-photon data (*Zhou et al., 2018)*), clustering based approaches (*Pachitariu et al., 2017*), dictionary learning (*Petersen et al., 2017*), or active contour models (*Reynolds et al., 2017*). Such spatio-temporal methods are unsupervised, and focus on detecting active neurons by considering the spatio-temporal activity of a component as a contiguous set of pixels within the FOV that are correlated in time. While such methods tend to offer a direct decomposition of the data in a set of sources with activity traces in an unsupervised way, in principle they require processing of the full dataset, and thus can be rendered intractable very quickly. Possible approaches to deal with the data size include distributed processing in High Performance Computing (HPC) clusters (*Freeman et al., 2014*), spatio-temporal decimation (*Friedrich et al., 2017a*), and dimensionality reduction (*Pachitariu et al., 2017*). Recently, *Giovannucci et al*. (*2017*) prototyped an online algorithm (OnACID), by adapting matrix factorization setups (*Pnevmatikakis et al., 2016; Mairal et al., 2010*), to operate on calcium imaging streaming data and thus natively deal with large data rates.

#### Deconvolution

For the problem of predicting spikes from fluorescence traces, both supervised and unsupervised methods have been explored. Supervised methods rely on the use of ground truth data to train or fit biophysical or neural network models (*Theis et al., 2016; Speiser et al., 2017*). Unsupervised methods can be either deterministic, such as sparse non-negative deconvolution (*Vogelstein et al., 2010; Pnevmatikakis et al., 2016)* that give a single estimate of the deconvolved neural activity, or probabilistic, that aim to also characterize the uncertainty around these estimates (e.g., (*Pnevmatikakis et al., 2013; Deneux et al., 2016*)). A recent community benchmarking effort (*Berens et al., 2017*) characterizes the similarities and differences of various available methods.

### CaImAn

Here we present CaImAn, an open source suite for the analysis pipeline of both two-photon and one-photon calcium imaging data. CaImAn includes frameworks for both offline analysis (CaImAn batch) where all the data is processed at once at the end of experiment, and online analysis on streaming data (CaImAn online). Moreover, CaImAn requires very moderate computing infrastructure (e.g., a personal laptop or workstation), thus providing automated, efficient, and reproducible large-scale analysis on commodity hardware.

## Contributions

Our contributions can be roughly grouped in three different directions:

**Methods:** CaImAn batch improves on the scalability of the source extraction problem by employing a MapReduce framework for parallel processing and memory mapping which allows the analysis of datasets larger than would fit in RAM on most computer systems. It also improves on the qualitative performance by introducing automated routines for component evaluation and classification, better handling of neuropil contamination, and better initialization methods. While these benefits are here presented in the context of the widely used CNMF algorithm of *Pnevmatikakis et al. (2016)*, they are in principle applicable to any matrix factorization approach. CaImAn online improves and extends the OnACID prototype algorithm (*Giovannucci et al., 2017*) by introducing, among other advances, new initialization methods and a convolutional neural network (CNN) based approach for detecting new neurons on streaming data. Our analysis on *in vivo* two-photon and light-sheet imaging datasets shows that CaImAn online approaches human-level performance and enables novel types of closed-loop experiments. Apart from these significant algorithmic improvements CaImAn includes several useful analysis tools such as, a MapReduce and memory-mapping compatible implementation of the CNMF-E algorithm for one-photon microendoscopic data (*Zhou et al., 2018*), a novel efficient algorithm for registration of components across multiple days, and routines for segmentation of structural (static) channel information which can be used for component seeding.
**Software:** CaImAn comes as a complete open source software suite implemented in Python, and is already widely used by, and has received contributions from, the community. It contains efficient implementations of the standard analysis pipeline steps (motion correction - source extraction - deconvolution - registration across different sessions), as well as numerous other features. Apart from Python, several of the tools presented here are also available in MATLAB^®^.
**Data:** We benchmark the performance of CaImAn against a previously unreleased corpus of manually annotated data. The corpus consists of 9 mouse *in vivo* two-photon datasets manually annotated by 3-4 independent labelers that were instructed to select active neurons in a principled and consistent way, and who subsequently combined their annotations to create a “consensus” ground truth that is also used to quantify the limits of human performance. The manual annotations are also released to the community providing a valuable tool for benchmarking and training purposes.

### Paper organization

The paper is organized as follows: We first give a brief presentation of the analysis methods and features provided by CaImAn. In the *Results* section we benchmark CaImAn online and CaImAn batch against a corpus of manually annotated data. We apply CaImAn online to a zebrafish whole brain lightsheet imaging recording, and demonstrate how such large datasets can be processed efficiently in real time. We also present applications of CaImAn batch to one-photon data, as well as examples of component registration across multiple days. We conclude by discussing the utility of our tools, the relationship between CaImAn batch and CaImAn online and outline future directions. Detailed descriptions of the introduced methods are presented in *Methods and Materials*.

## Methods

Before presenting the new analysis features introduced with this work, we overview the analysis pipeline that CaImAn uses and builds upon.

### Overview of analysis pipeline

The standard analysis pipeline for calcium imaging data used in CaImAn is depicted in Fig. 1a. The data in movie format is first processed to remove motion artifacts. Subsequently the active components (neurons and background) are extracted as individual pairs of a spatial footprint that describes the shape of each component projected to the imaged FOV, and a temporal trace that captures its fluorescence activity (Fig. 1 b-d). Finally, the neural activity of each fluorescence trace is deconvolved from the dynamics of the calcium indicator. These operations can be challenging because of limited axial resolution of 2-photon microscopy (or the much larger integration volume in one-photon imaging). This results in spatially overlapping fluorescence from different sources and neuropil activity. Before presenting the new features of CaImAn in more detail, we briefly review how it incorporates existing tools in the pipeline.

#### Motion Correction

CaImAn uses the NoRMCorre algorithm (*Pnevmatikakis and Giovannucci, 2017*) that corrects non-rigid motion artifacts by estimating motion vectors with subpixel resolution over a set of overlapping patches within the FOV. These estimates are used to infer a smooth motion field within the FOV for each frame. For two-photon imaging data this approach is directly applicable, whereas for one-photon micro-endoscopic data the motion is estimated on high pass spatially filtered data, a necessary operation to remove the smooth background signal and create enhanced spatial landmarks. The inferred motion fields are then applied to the original data frames.

#### Source Extraction

Source extraction is performed using the constrained non-negative matrix factorization (CNMF) framework of *Pnevmatikakis et al. (2016)* which can extract components with spatial overlapping projections (Fig. 1b). After motion correction the spatio-temporal activity of each source can be expressed as a rank one matrix given by the outer product of two components: a component in space that describes the spatial footprint (location and shape) of each source, and a component in time that describes the activity trace of the source (Fig. 1 c). The data can be described by the sum of all the resulting rank one matrices together with an appropriate term for the background and neuropil signal and a noise term (Fig. 1d). For two-photon data the neuropil signal can be modeled as a low rank matrix (*Pnevmatikakis et al., 2016*). For microendoscopic data the larger integration volume leads to more complex background contamination (*Zhou et al., 2018*). Therefore, a more descriptive model is required (see *Methods and Materials (Mathematical model of the CNMF framework*) for a mathematical description). CaImAn batch embeds these approaches into a general algorithmic framework that enables scalable automated processing with improved results in terms of quality and processing speed.

#### Deconvolution

Neural activity deconvolution is performed using sparse non-negative deconvolution (*Vogelstein et al., 2010; Pnevmatikakis et al., 2016*) and implemented with both the near-online OASIS algorithm (*Friedrich et al., 2017b*) and an efficient convex optimization framework (*Pnevmatikakis et al., 2016*). The algorithm is competitive to the state of the art according to recent benchmarking studies (*Berens et al., 2017*). Prior to deconvolution, the traces are detrended to remove non-stationary effects, e.g., photo-bleaching.

#### Online Processing

The three processing steps described above can be implemented in an online fashion on streaming data using the OnACID algorithm (*Giovannucci et al., 2017*). The method builds upon the online dictionary learning framework presented in *Mairal et al*. (*2010*) for source extraction, by adding the capability of finding new components as they appear and also incorporating the steps of motion correction and deconvolution (Fig. 1e). CaImAn online extends and improves the OnACID prototype algorithm by introducing a number of algorithmic features and a CNN based component detection approach, leading to a major performance improvement.

We now present the new methods introduced by CaImAn. More details are given in *Methods and Materials* and pseudocode descriptions of the main routines are given in the *Appendix*.

### Batch processing of large scale datasets on standalone machines

The batch processing pipeline mentioned above can become a computational bottleneck when tackled without customized solutions. For instance, a naive approach to the problem might have as a first step to load in-memory the full dataset; this approach is non-scalable as datasets typically exceed available RAM (and extra memory is required by any analysis pipeline). To limit memory usage, as well as computation time, CaImAn batch relies on a MapReduce approach (*Dean and Ghemawat, 2008*). Unlike previous work (*Freeman et al., 2014*), CaImAn batch assumes minimal computational infrastructure (up to a standard laptop computer), is not tied to a particular parallel computation framework, and is compatible with HPC scheduling systems like SLURM (*Yoo et al., 2003*).

Naive implementations of motion correction algorithms need to either load in memory the full dataset or are constrained to process one frame at a time, therefore preventing parallelization. Motion correction is parallelized in CaImAn batch without significant memory overhead by processing several temporal chunks of a video data on different CPUs. CaImAn batch broadcasts to each CPU a meta-template, which is used to align all the frames in the chunk. Each process writes in parallel to the target file containing motion-corrected data, which is stored in as a memory mapped array. This allows arithmetic operations to be performed against data stored on the hard drive with minimal memory use, and slices of data to be indexed and accessed without loading the full file in memory. More details are given in *Methods and Materials (Memory mapping*).

Similarly, the source extraction problem, especially in the case of detecting cell bodies, is inherently local with a neuron typically appearing in a neighborhood within a small radius from its center of mass (Fig. 2a). Exploiting this locality, CaImAn batch splits the FOV into a set of spatially overlapping patches which enables the parallelization of the CNMF (or any other) algorithm to extract the corresponding set of local spatial and temporal components. The user specifies the size of the patch, the amount of overlap between neighboring patches and the initialization parameters for each patch (number of components and rank background for CNMF, stopping criteria for CNMF-E). Subsequently the patches are processed in parallel by the CNMF/CNMF-E algorithm to extract the components and neuropil signals from each patch.

**Figure 2.**
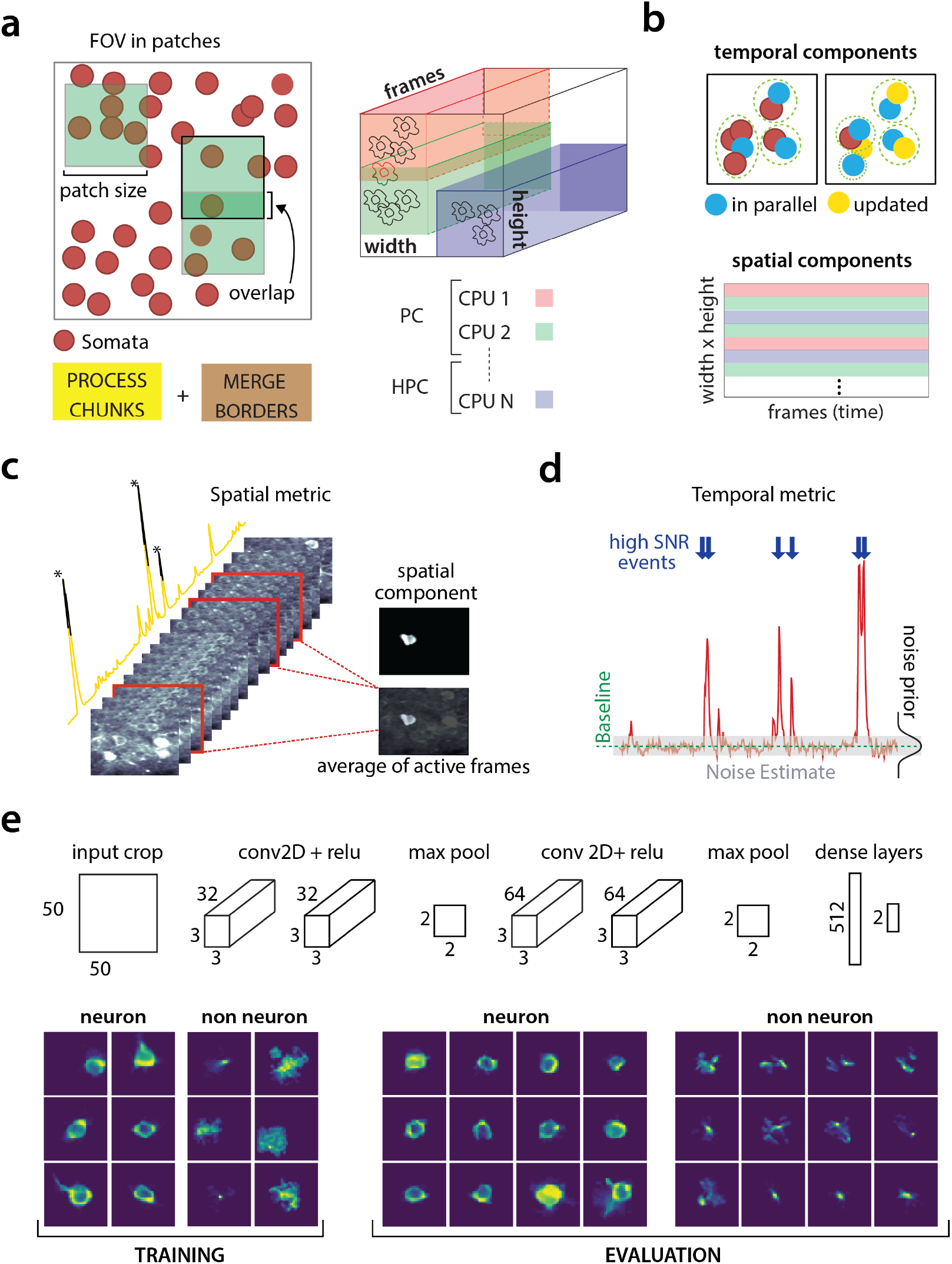
Parallelized processing and component quality assessment for CaImAn batch. (a) Illustration of the parallelization approach used by CaImAn batch for source extraction. The data movie is partitioned into overlapping sub-tensors, each of which is processed in an embarrassingly parallel fashion using CNMF, either on local cores or across several machines in a HPC. The results are then combined. (b) Refinement after combining the results can also be parallelized both in space and in time. Temporal traces of spatially non-overlapping components can be updated in parallel (top) and the contribution of the spatial footprints for each pixel can be computed in parallel (bottom). Parallelization in combination with memory mapping enable large scale processing with moderate computing infrastructure. (c) Quality assessment in space: The spatial footprint of each real component is correlated with the data averaged overtime, after removal of all other activity. (d) Quality assessment in time: A high SNR is typically maintained over the course of a calcium transient. (e) CNN based assessment. *Top:* A 4-layer CNN based classifier is used to classify the spatial footprint of each component into neurons or not. *Bottom:* Positive and negative examples for the CNN classifier, during training (left) and evaluation (right) phase. The CNN classifier can accurately classify shapes and generalizes across datasets from different brain areas.

Apart from harnessing memory and computational benefits due to parallelization, processing in patches acts indirectly as a dynamic range equalizer and enables CaImAn batch to detect neurons across the whole FOV, a feature absent in the original CNMF, where areas with high absolute fluorescence variation tend to be favored. This results in better source extraction performance. After all the patches have been processed, the results are embedded within the FOV (Fig. 2a), and the overlapping regions between neighboring patches are processed so that components corresponding to the same neuron are merged. The process is summarized in algorithmic format in Alg. 1 and more details are given in *Methods and Materials (Combining results from different patches*).

### Initialization Methods

Initialization methods for matrix factorization problems can impact results due to the non-convex nature of their objective function. CaImAn batch provides an extension of the GreedyROI method used in *Pnevmatikakis et al. (2016)*, that detects neurons based on localized spatiotemporal activity. CaImAn batch can also be seeded with binary masks that are obtained from different sources, e.g., through manual annotation or segmentation of structural channel (SeededInitialization, Alg. 2). More details are given in *Methods and Materials (Initialization strategies*).

### Automated component evaluation and classif cation

A common limitation of matrix factorization algorithms is that the number of components that the algorithm seeks during its initialization must be pre-determined by the user. For example, *Pnevmatikakis et al*. (*2016*) suggest a large number of components which are then heuristically ordered according to their size and activity pattern. When processing large datasets in patches the target number of components is passed on to every patch implicitly assuming a uniform density of (active) neurons within the entire FOV. In general this assumption does not hold and can generate a large number of spurious components. CaImAn introduces tests to assess the quality of the detected components and eliminate possible false positives. These tests are based on the observation that active components are bound to have a distinct localized spatio-temporal signature within the FOV. We present below unsupervised and supervised tests employed by CaImAn for component classification. In CaImAn batch, they are initially applied after the processing of each patch is completed, and additionally as a post-processing step after the results from the patches have been merged and refined, whereas in CaImAn online they are used to screen new candidate components. We briefly present these tests below and refer to *Methods and Materials (Details of quality assessment tests)* for more details:

**Spatial footprint consistency:** To test whether a detected component is spurious, we correlate the spatial footprint of this component with the average frame of the data, taken over the interval when the component, with no other overlapping component, was active (Fig. 2c). The component is rejected if the correlation coefficient is below a certain threshold *θ*_sp_ (e.g., *θ*_sp_ < 0.05).
**Trace SNR:** Similarly, for each component we computed the peak SNR of its temporal trace averaged over the duration of a typical transient (Fig. 2d). The component is rejected if the computed SNR is below a certain threshold *θ*_SNR_ (e.g., *θ*_SNR_ = 2).
**CNN based classification:** We also trained a 4-layer convolutional neural network (CNN) to classify spatial footprints into true or false components (Fig. 2e), where a true component here corresponds to a spatial footprint that resembles the soma of a neuron. The classifier, named batch classifier, was trained on a small corpus of manually annotated datasets (full description given in section *Benchmarking against ground truth*) and exhibited similar high classification performance on test samples from different datasets.

While CaImAn uses the CNMF algorithm, the tests described above can be applied to results obtained from any source extraction algorithm, highlighting the modularity of our tools.

### Online analysis with CaImAn online

CaImAn supports online analysis on streaming data building on the core of the prototype algorithm of *Giovannucci et al*. (*2017*), and extending it in terms of qualitative performance and computational efficiency:

**Initialization:** Apart from initializing CaImAn online with CaImAn batch on a small time interval, CaImAn online can also be initialized in a bare form over an even smaller time interval, where only the background components are estimated and all the components are determined during the online analysis. This process, named BareInitialization, can be achieved by running the CNMF algorithm (*Pnevmatikakis et al., 2016*) over the small interval to estimate the background components and possibly a small number of components. The SeededInitialization of Alg. 2 can also be used.
**Deconvolution:** Instead of a separate step after demixing as in *Giovannucci et al*. (*2017*), deconvolution here can be performed simultaneously with the demixing online, leading to more stable traces especially in cases of low-SNR, as also observed in *Pnevmatikakis et al*. (*2016*). Online deconvolution can also be performed for models that assume second order calcium dynamics, bringing the full power of *Friedrich et al. (2017b)* to processing of streaming data.
**Epochs:** CaImAn online supports multiple passes over the data, a process that can detect early activity of neurons that were not picked up during the initial pass, as well as smooth the activity of components that were detected at late stages during the first epoch.
**New component detection using a CNN:** To search for new components in a streaming setup, OnACID keeps a buffer of the residual frames, computed by subtracting the activity of already found components and background signals. Candidate components are determined by looking for points of maximum energy in this residual signal, after some smoothing and dynamic range equalization. For each such point identified, a candidate shape and trace are constructed using a rank-1 NMF in a local neighborhood around this point. In its original formulation (*Giovannucci et al., 2017*), the shape of the component was evaluated using the space correlation test described above. Here, we introduce a CNN classifier approach that tests candidate components by examining their spatial footprint as obtained by the average of the residual buffer across time. This online classifier (different from the batch classifier for quality assessment described above), is trained to be strict, minimizing the number of false positive components that enter the online processing pipeline. It can test multiple components in parallel, and it achieves better performance with no hyper-parameter tuning compared to the previous approach. More details on the architecture and training procedure are given in *Methods and Materials (Classification through CNNs)*. The identification of candidate components is further improved by performing spatial high pass filtering on the average residual buffer to enhance its contrast. The new process for detecting neurons is described in Algs. 3 and 4. See Supplemental Movies 1 and 2 on a detailed graphic description of the new component detection step.

### Component registration across multiple sessions

CaImAn provides a method to register components from the same FOV across different sessions. The method uses a simple intersection over union metric to calculate the distance between different cells in different sessions and solving a linear assignment problem to perform the registration in a fully automated way (RegisterPair, Alg. 5). To register the components between more than 2 sessions (RegisterMulti, Alg. 6), we order the sessions chronologically and register the components of the current session against the union of component of all the past sessions aligned to the current FOV. This allows for the tracking of components across multiple sessions without the need of pairwise registration between each pair of sessions. More details as well as discussion of other methods (*Sheintuch et al., 2017)* are given in *Methods and Materials (Component registration)*.

### Benchmarking against ground truth

To quantitatively evaluate CaImAn we benchmarked its results against ground truth data.

#### Creating ground truth data through manual annotation

We collected manual annotations from multiple independent labelers who were instructed to find round or donut shaped^2^ *active* neurons on 9 two-photon *in vivo* mouse brain datasets. The datasets were collected at various labs and from various brain areas (hippocampus, visual cortex, parietal cortex) using several GCaMP variants. A summary of the features of all the annotated datasets is given in Table 2. Details about the annotation procedure are given in *Methods and Materials*.

To address human variability in manual annotation each dataset was labeled by 3 or 4 independent labelers, and the final ground truth dataset was created by having the different labelers reaching a *consensus* over their disagreements (Fig. 3a). The result of this process was defined as ground truth for the evaluation of CaImAn as well as each individual labeler against the consensus (Fig. 3b)^3^. More details are given in *Methods and Materials (Collection of manual annotations and*

**Figure 3.**
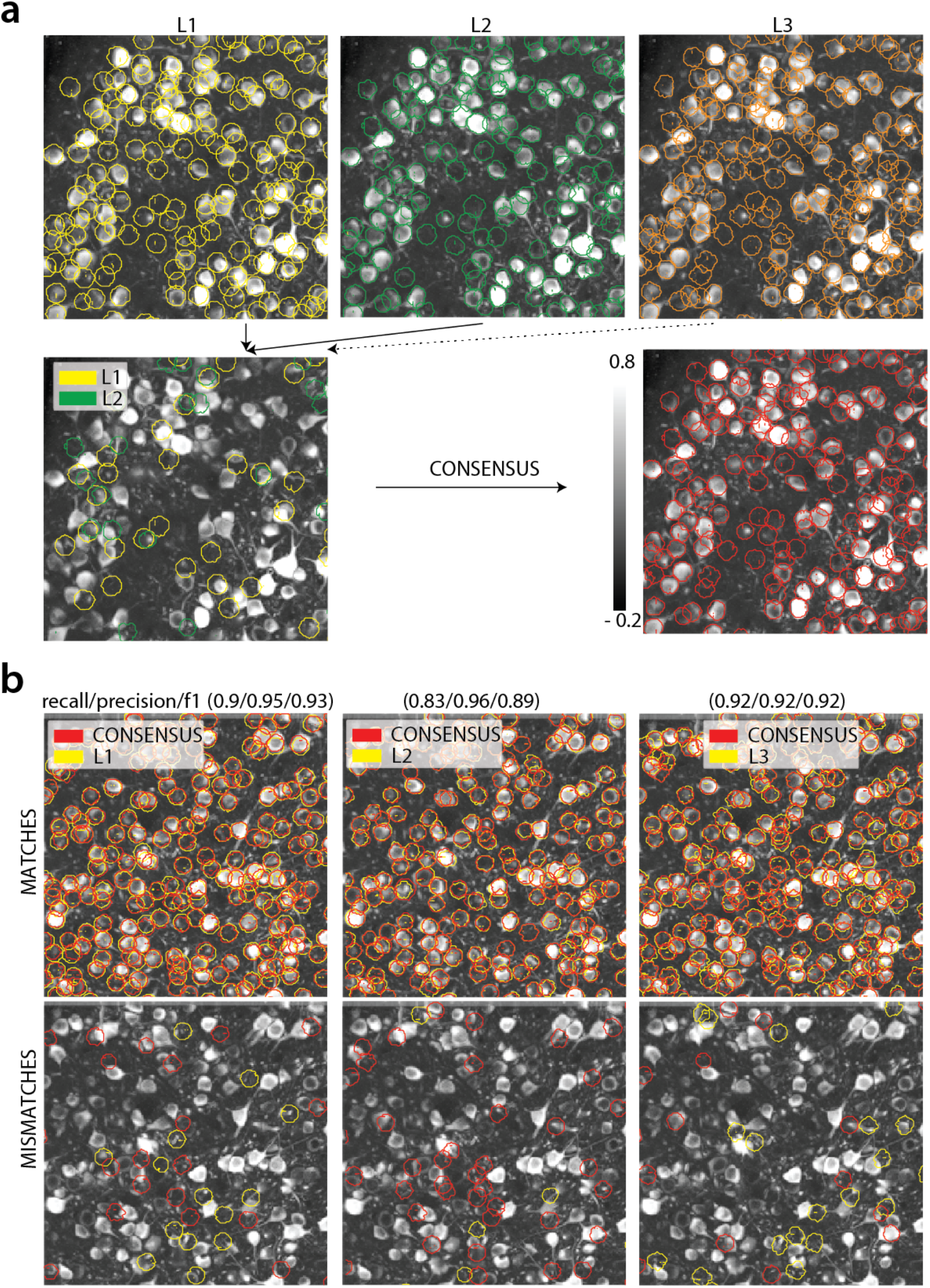
Ground truth generation. (a) *Top:* Individual manual annotations on the dataset N.04.00.t (only part of the FOV is shown) for labelers L1 (left), L2 (middle), L3(right). *Bottom:* Disagreements between L1 and L2 (left), and ground truth labels (right) after the consensus between all labelers has been reached. In this example, consensus considerably reduced the number of initially selected neurons. (b) Matches (top) and mismatches (bottom) between each individual labeler and consensus ground truth. Red contours on the mismatches panels denote false negative contours, i.e., components in the consensus not selected by the corresponding labeler, whereas yellow contours indicate false positive contours. Performance of each labeler is given in terms of precision/recall and *F*_1_ score and indicates an unexpected level of variability between individual labelers.

*ground truth*). We believe that the current database, which will be made publicly available, presents an improvement over the existing neurofinder database (http://neurofinder.codeneuro.org/) in several aspects:

**Consistency:** The datasets are annotated using exactly the same procedure (see *Methods and Materials*), and in all datasets the goal is to detect only active cells. In contrast, the annotation of the various neurofinder datasets is performed either manually or automatically by segmenting an image of a static (structural) indicator. Even though structural indicators could be used for ground truth extraction, the segmentation of such images is not a straightforward problem in the case of dense expression, and the stochastic expression of indicators can lead to mismatches between functional and structural indicators.
**Uncertainty quantification:** By employing more than one human labeler we discovered a surprising level of disagreement between different annotators (see Table 1, Fig. 3b for details), which renders individual annotations somewhat unreliable for benchmarking purposes, and non-reproducible. The combination of the various annotations leads to more reliable ground truth and also quantifies the limits of human performance.

**Table 1.** Results of each labeler, CaImAn batch and CaImAn online algorithms against consensus ground truth. Results are given in the form *F*_1_ score (precision, recall), and empty entries correspond to datasets not manually annotated by the specific labeler. In *italics* the datasets used to train the CNN classifiers.

| Name | L1 | L2 | L3 | L4 | CALMAN BATCH | CALMAN ONLINE |
| --- | --- | --- | --- | --- | --- | --- |
| *N. 03. 00. t* | X | 0.90<br>(0.88,0.92) | 0.85<br>(0.78,0.93) | 0.78<br>(0.73,0.83) | 0.78<br>(0.77,0.79) | 0.76<br>(0.77,0.75) |
| *N. 04. 00. t* | X | 0.69<br>(0.54,0.97) | 0.75<br>(0.61,0.97) | 0.87<br>(0.78,0.98) | 0.67<br>(0.62,0.72) | 0.68<br>(0.65,0.71) |
| *N. 02. 00* | 0.89<br>(0.86,0.93) | 0.87<br>(0.88,0.85) | 0.84<br>(0.92,0.77) | 0.82<br>(1.00,0.70) | 0.79<br>(0.8,0.77) | 0.77<br>(0.79,0.76) |
| *N. 00. 00* | X | 0.92<br>(0.93,0.91) | 0.83<br>(0.86,0.80) | 0.87<br>(0.96,0.80) | 0.72<br>(0.83,0.64) | 0.72<br>(0.83,0.64) |
| *N. 01. 01* | 0.80<br>(0.95,0.69) | 0.89<br>(0.96,0.83) | 0.78<br>(0.73,0.84) | 0.75<br>(0.80,0.70) | 0.77<br>(0.88,0.69) | 0.73<br>(0.78,0.68) |
| YST | 0.78<br>(0.76,0.81) | 0.90<br>(0.85,0.97) | 0.82<br>(0.75,0.92) | 0.79<br>(0.96,0.67) | 0.76<br>(0.9,0.66) | 0.78<br>(0.76,0.81) |
| K53 | 0.89<br>(0.96,0.83) | 0.92<br>(0.92,0.92) | 0.93<br>(0.95,0.91) | 0.83<br>(1.00,0.72) | 0.77<br>(0.83,0.72) | 0.82<br>(0.80,0.83) |
| J115 | X | 0.93<br>(0.94,0.91) | 0.94<br>(0.95,0.93) | 0.83<br>(1.00,0.71) | 0.77<br>(0.9,0.68) | 0.81<br>(0.75,0.88) |
| J123 | 0.85<br>(0.96,0.76) | 0.83<br>(0.73,0.96) | 0.90<br>(0.91,0.90) | 0.91<br>(0.92,0.89) | 0.68<br>(0.94,0.51) | 0.80<br>(0.82,0.79) |

#### Comparing CaImAn against ground truth

To compare CaImAn against the consensus ground truth, the manual annotations were used as binary masks to construct the ground truth spatial and temporal components, using the SeededInitialization procedure (Alg. 2) of CaImAn batch. The set of spatial footprints obtained from CaImAn is registered against the set of ground truth spatial footprints (derived as described above) using the RegisterPair algorithm (Alg. 5) for component registration described above. Performance is then quantified using a precision/recall framework similar to other studies (*Apthorpe et al., 2016; Giovannucci et al., 2017*).

### Software

CaImAn is developed by and for the community. Python open source code for all the methods described above is available at https://github.com/flatironinstitute/CaImAn. The repository contains documentation, numerous demos, and Jupyter notebook tutorials, as well as visualization tools, and a message/discussion board. The code, which is compatible with Python 2 and Python 3^4^, uses tools from several open source libraries, such as OpenCV (*Bradski, 2000*), scikit-learn (*Pedregosa et al., 2011*), and scikit-image (*Van der Walt et al., 2014*). Most routines are also available in MATLAB^®^ at https://github.com/flatironinstitute/CaImAn-MATLAB.

## Results

### Manual annotations show a high degree of variability

We compared the performance of each human annotator against a consensus ground truth. The performance was quantified with a precision/recall framework and the results of the performance of each individual labeler against the consensus ground truth for each dataset is given in Table 1. The range of human performance in terms of *F*_1_ score was 0.69-0.94, with average 0.83± 0.07 (mean ± STD). All annotators performed similarly on average (0.83±0.05, 0.83±0.08, 0.84±0.06, 0.85±0.08). We also ensured that the performance of labelers was stable across time (i.e. their learning curve plateaued, data not shown). As shown in Table 1 (see also Fig 4b) the *F*_1_ score was never 1, and in most cases it was less or equal to 0.9, demonstrating significant variability between annotators. Fig. 3 (bottom) shows an example of matches and mismatches between individual labelers and consensus ground truth for dataset K53, where the level of agreement was relatively high. The high degree of variability in human responses indicates the challenging nature of the source extraction problem and raises reproducibility concerns in studies relying heavily on manual ROI selection.

### CaImAn batch and CaImAn online detect neurons with near-human accuracy

We first benchmarked CaImAn batch and CaImAn online against consensus ground truth for the task of identifying neurons locations and their spatial footprints, using the same precision recall framework (Table 1). Fig. 4a shows an example dataset (K53) along with neuron-wise matches and mismatches between CaImAn batch and consensus ground truth (top) and CaImAn online vs consensus ground truth (bottom).

**Figure 4.**
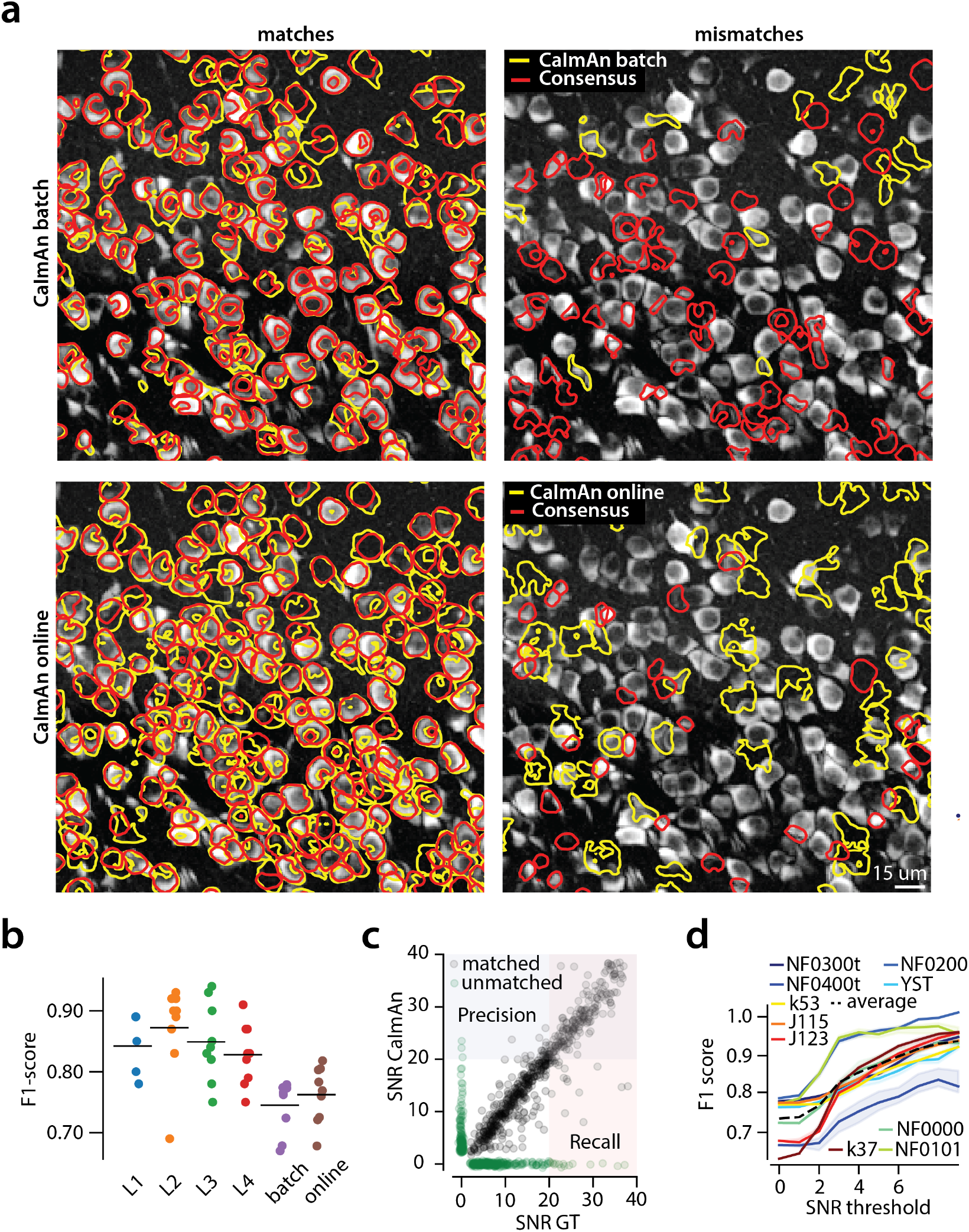
Evaluation of CaImAn performance against manually annotated data. (a) Comparison of CaImAn batch (top) and CaImAn online (bottom) when benchmarked against consensus ground truth for dataset K53. Fora portion of the FOV, correlation image overlaid with matches (left panels, true positives red for consensus ground truth, yellow for CaImAn) and mismatches (right panels, red for false negatives, yellow for false positives). (b) Performance of CaImAn batch, CaImAn online and all labelers (L1, L2, L3, L4) for all 9 datasets in terms of *F_1_* score. CaImAn batch and CaImAn online reach near-human accuracy for neuron detection. Complete results with precision and recall for each dataset are given in Table 1. (c-d) Performance of CaImAn batch increases with peak SNR. (c) Example of scatter plot between SNRs of matched traces between CaImAn batch and ground truth for dataset K53. False negative/positive pairs are plotted in green along the x- and y-axes respectively, perturbed as a point cloud to illustrate the density. Most false positive/negative predictions occur at low SNR values. Shaded areas represent thresholds above which components are considered for matching (blue for CaImAn batch selected components and red for GT selected components) (d) *F_1_* score and upper/lower bounds for all datasets as a function of various peak SNR thresholds. Performance increases significantly for neurons with high peak SNR traces (see text for definition of metrics and the bounds).

The results indicate a similar performance between CaImAn batch and CaImAn online; CaImAn batch has *F*_1_ scores in the range 0.68-0.79 and average performance 0.75±0.04 (mean±STD). On the other hand CaImAn online had *F*_1_ scores in the range 0.68-0.82 and average performance 0.76±0.04. While the two algorithms performed similarly on average, CaImAn batch tends to perform better for shorter datasets whereas online processing tends to lead to better results for longer datasets (see Table 2 for characteristics of the various datasets). CaImAn approaches but is in most cases below the accuracy levels of human annotators (Fig. 4b). This can be attributed to a number of reasons: First, to demonstrate the generality and ease of use of our tools, the results presented here are obtained by running CaImAn batch and CaImAn online with *exactly* the same parameters for each dataset (see *Methods and Materials (Implementation details)*): fine-tuning to each individual dataset can significantly increase performance. Second, CNMF detects active components regardless of their shape, and can detect non-somatic structures with significant transients. While non-somatic components can be filtered out to some extent using the CNN classifier, their existence degrades performance compared to the ground truth that consists only of neurons. Lastly, the ground truth is by construction a subset of the union of all individual annotations, which can bias upwards the scores of individual labelers.

**Table 2.**
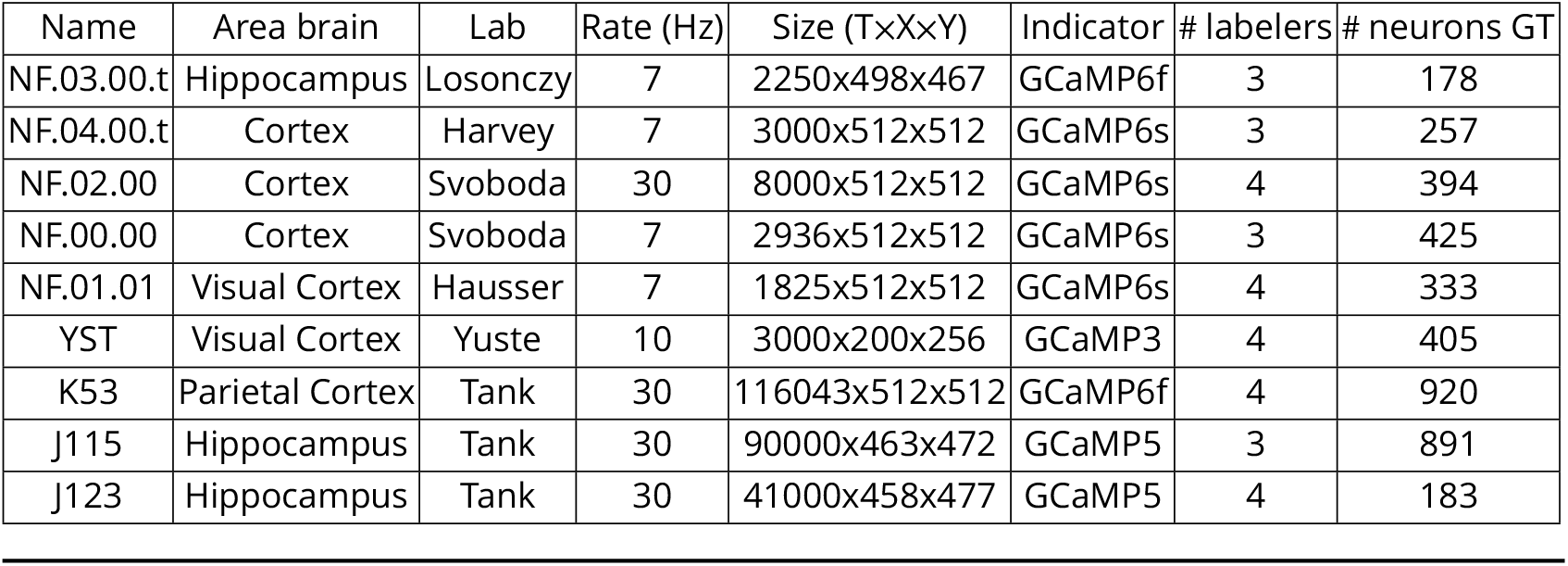
Properties of manually annotated datasets. For each dataset the duration, imaging rate and calcium indicator are given, as well as the number of active neurons selected after consensus of the manual annotations.

### Neurons with higher SNR transients are detected more accurately

While CaImAn online had balanced performance with respect to precision and recall (mean precision 0.77±0.05, mean recall 0.76±0.07), CaImAn batch showed significantly higher precision than recall (mean precision 0.83±0.09, mean recall 0.69±0.08). We looked into this behavior, by analyzing CaImAn batch performance as a function of the SNR of the inferred and ground truth traces (Fig. 4c-d). The SNR measure of a trace corresponds to the peak-SNR averaged over the length of a typical trace (see *Methods and Materials (Detecting fluorescence traces with high SNR)*). An example is shown in Fig. 4c where the scatter plot of SNR between matched ground truth and inferred traces is shown (false negative/positive components are shown along the x- and y- axis, respectively). To evaluate the performance we computed a precision metric as the fraction of inferred components above a certain SNR threshold that are matched with a ground truth component (Fig. 4c, shaded blue). Similarly we computed a recall metric as the fraction of ground truth components above a SNR threshold that are detected by CaImAn batch (Fig. 4c, shaded red), and an *F*_1_ score as the harmonic mean of the two (Fig. 4d). The results indicate that the performance significantly grows as a function of the SNR for all datasets considered, growing on average from 0.73 when all neurons are considered to 0.92 when only neurons with traces having SNR ≥ 9 are considered (Fig. 4d)^5^.

### CaImAn reproduces the ground truth traces with high fidelity

Testing the quality of the inferred traces is a more challenging task due to the complete lack of ground truth data in the context of large scale *in vivo* recordings. As mentioned above, we considered as ground truth the traces obtained by running the CNMF algorithm seeded with the binary masks obtained by consensus ground truth procedure. After alignment of the ground truth with the results of CaImAn, the matched traces were compared both for CaImAn batch and for CaImAn online. Fig. 5a, shows an example of 5 of these traces for the dataset K53, showing very similar behavior of the traces in these three different cases.

**Figure 5.**
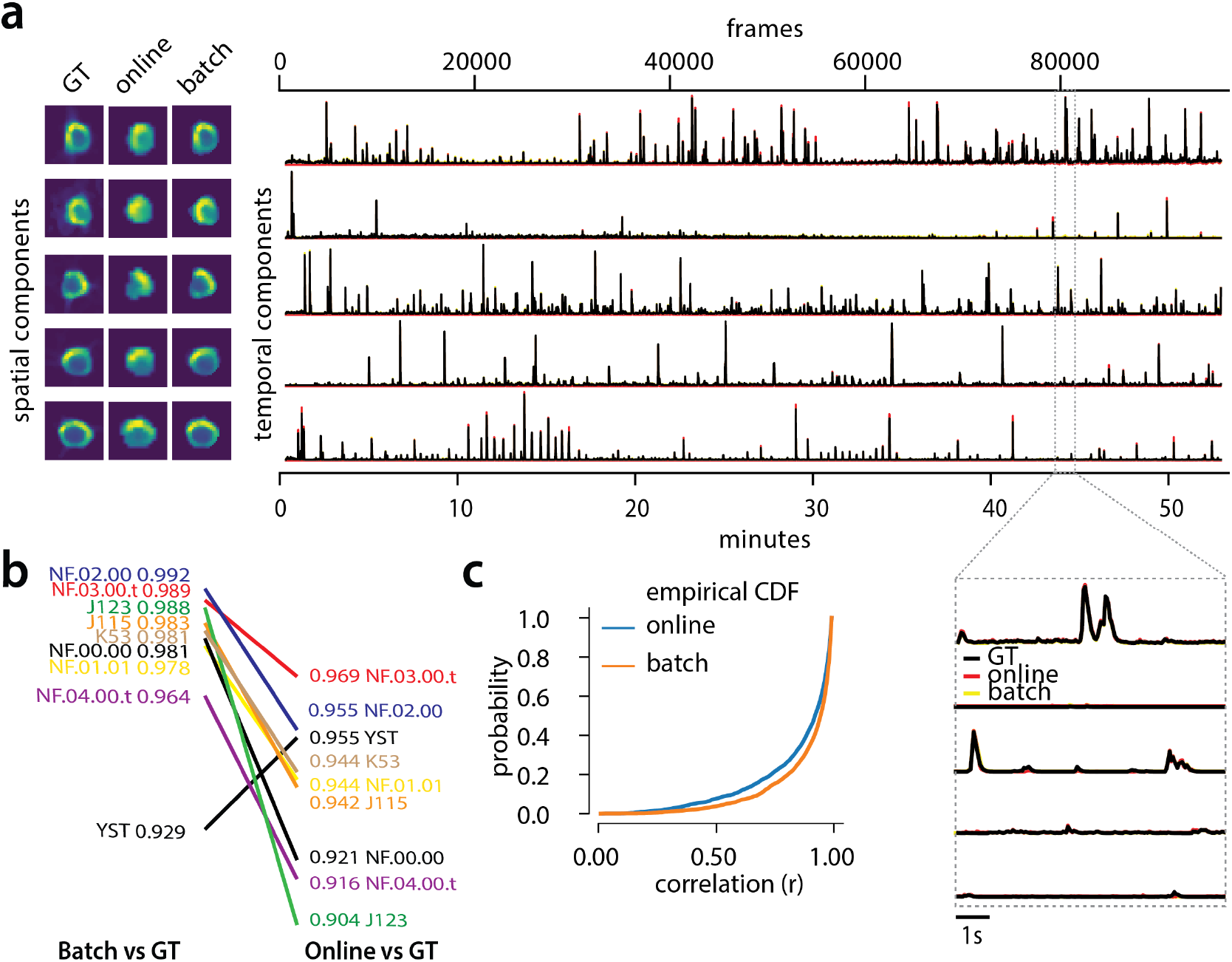
Evaluation of CaImAn extracted traces against traces derived from ground truth. (a) Examples of shapes (left) and traces (right) are shown for five matched components extracted from dataset K53 for consensus ground truth (GT, black), CaImAn batch (yellow) and CaImAn online (red) algorithms. The dashed gray portion of the traces is also shown magnified (bottom-right). Spatial footprints and traces for ground truth are obtained by seeding CaImAn with the consensus binary masks. The traces extracted from both versions of CaImAn match closely the ground truth traces. (b) Slope graph for the average correlation coefficient for matches between ground truth and CaImAn batch, and between ground truth and CaImAn online. Batch processing produces traces that match more closely the traces extracted from the ground truth data. (c) Empirical cumulative distribution functions of correlation coefficients aggregated over all the tested datasets. Both distributions exhibit a sharp derivative close 1 (last bin), with the batch approach giving better results.

To quantify the similarity we computed the correlation coefficients of the traces (ground truth vs CaImAn batch, and ground truth vs CaImAn online) for all the 9 datasets (Fig. 5b-c). Results indicated that for all but one dataset (Fig. 5b) CaImAn batch reproduced the traces with higher fidelity, and in all cases the mean correlation coefficients was higher than 0.9, and the empirical histogram of correlation coefficients peaked at the maximum bin 0.99-1 (Fig. 5c). The results indicate that the batch approach extracts traces closer to the ground truth traces. This can be attributed to a number of reasons: By processing all the time points simultaneously, the batch approach can smooth the trace estimation over the entire time interval as opposed to the online approach where at each timestep only the information up to that point is considered. Moreover, CaImAn online might not detect a neuron until it becomes strongly active. This neuron’s activity before detection is unknown and has a default value of zero, resulting in a lower correlation coefficient. While this can be ameliorated to a great extent with additional passes over the data, the results indicate trade-offs between using the online and offline versions of CaImAn.

### Online analysis of a whole brain zebrafish dataset

We tested CaImAn online with a 380GB whole brain dataset of larval zebrafish (*Danio rerio*) acquired with a light-sheet microscope (*Kawashima et al., 2016*). The imaged transgenic fish (Tg(elavl3:H2B-GCaMP6f)jf7) expressed the genetically encoded calcium indicator GCaMP6f in almost all neuronal nuclei. Data from 45 planes (FOV 820×410 *μ*m^2^, spaced at 5.5 *μ*m intervals along the dorso-ventral axis) was collected at 1Hz for 30 minutes (for details about preparation, equipment and experiment refer to *Kawashima et al. (2016)*). With the goal of simulating real-time analysis of the data, we run all the 45 planes in parallel on a computing cluster with 9 nodes (each node is equipped with 24 CPUs and 128-256 GB RAM). Data was not stored locally in each machine but directly accessed from a network drive.

The algorithm was initialized with CaImAn batch run on 200 initial frames and looking for 500 components. The small number of frames (1885) and the large FOV size (2048 × 1188 pixels) for this dataset motivated this choice of increased number of components during initialization. In Fig. 6 we report the results of the analysis for plane number 11 of 45. For plane 11, CaImAn online found 1524 neurons after processing 1685 frames. Since no ground truth was available for this dataset, it was only possible to evaluate the performance of this algorithm by visual inspection. CaImAn online identified all the neurons with a clear footprint in the underlying correlation image (higher SNR, Fig. 6a) and missed a small number of the fainter ones (low SNR). By visual inspection of the components the authors could find very few false positives. Given that the parameters were not tuned and that the classifier was not trained on zebrafish neurons, we hypothesize that the algorithm is biased towards a high precision result. Spatial components displayed the expected morphological features of neurons (Fig. 6b-c). Considering all the planes (Figs 6e and 11) CaImAn online was able to identify in a single pass of the data a total of 66108 neurons. See Supplemental Movie 3 for a summary across all planes. The analysis was performed in 21 minutes, with the first 3 minutes allocated to the initialization and the remaining 18 to process the rest of the data in streaming mode (and in parallel for each plane). This demonstrates the ability of CaImAn online to process large amounts of data in real-time (see also Fig. 8 for a discussion of computational performance).

**Figure 6.**
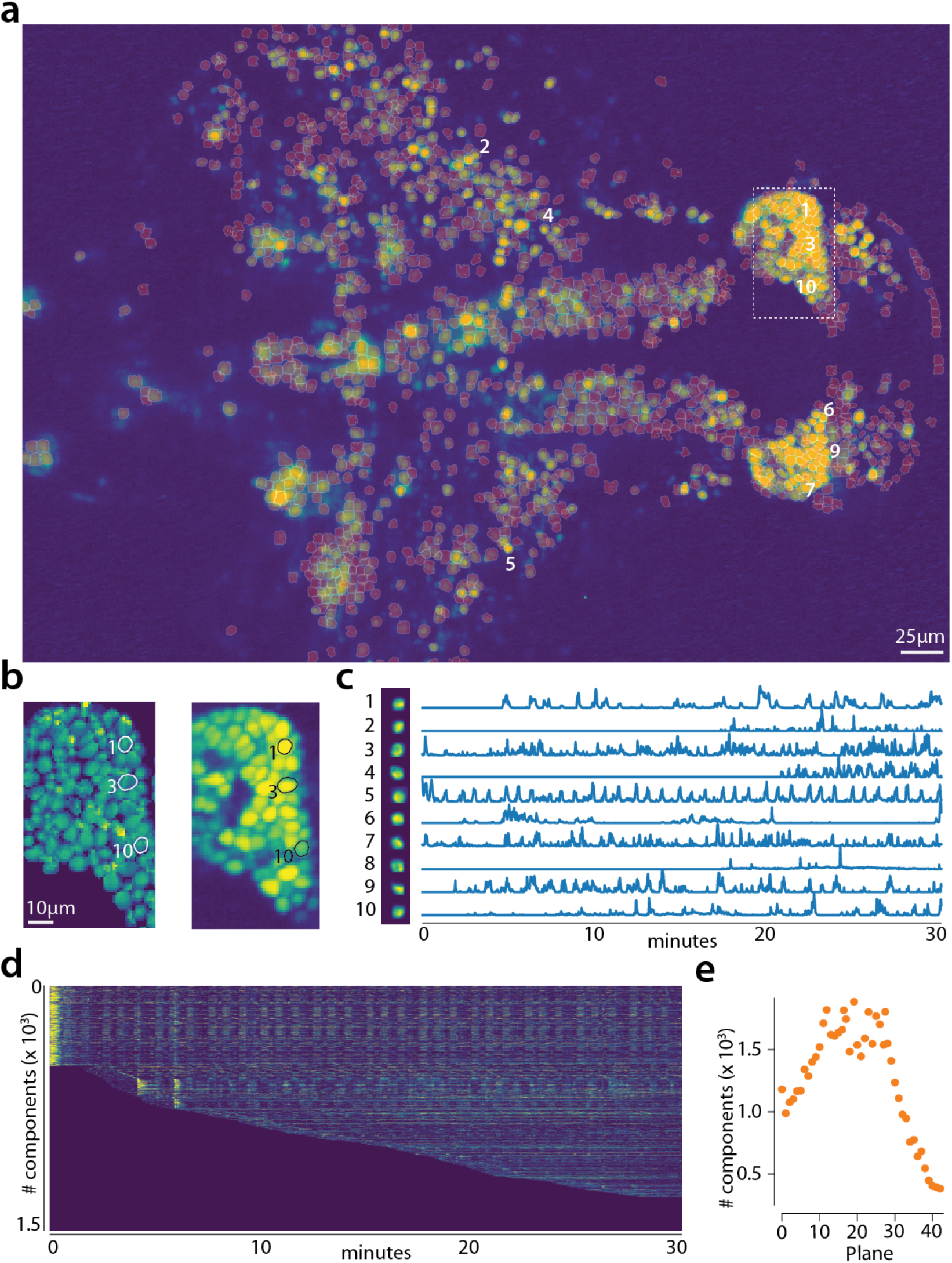
Online analysis of a 30 min long whole brain recording of the zebraf¡sh brain. (a) Correlation image overlaid with the spatial components found by the algorithm (portion of plane 11 out of 45 planes in total). (b) Left: Spatial footprints found in the dashed region in (a), contours represent neurons displayed in (c). Right: Correlation image for the same region. (c) Spatial (left) and Temporal (right) components associated to the ten example neurons marked in panel (a). (d) Temporal traces for all the neurons found in the FOV in (a), the initialization on the first 200 frames contained 500 neurons (present since time 0). (e) Number of neurons found per plane (See also Supplementary Fig. 11 for a summary of the results from all planes).

**Figure 7.**
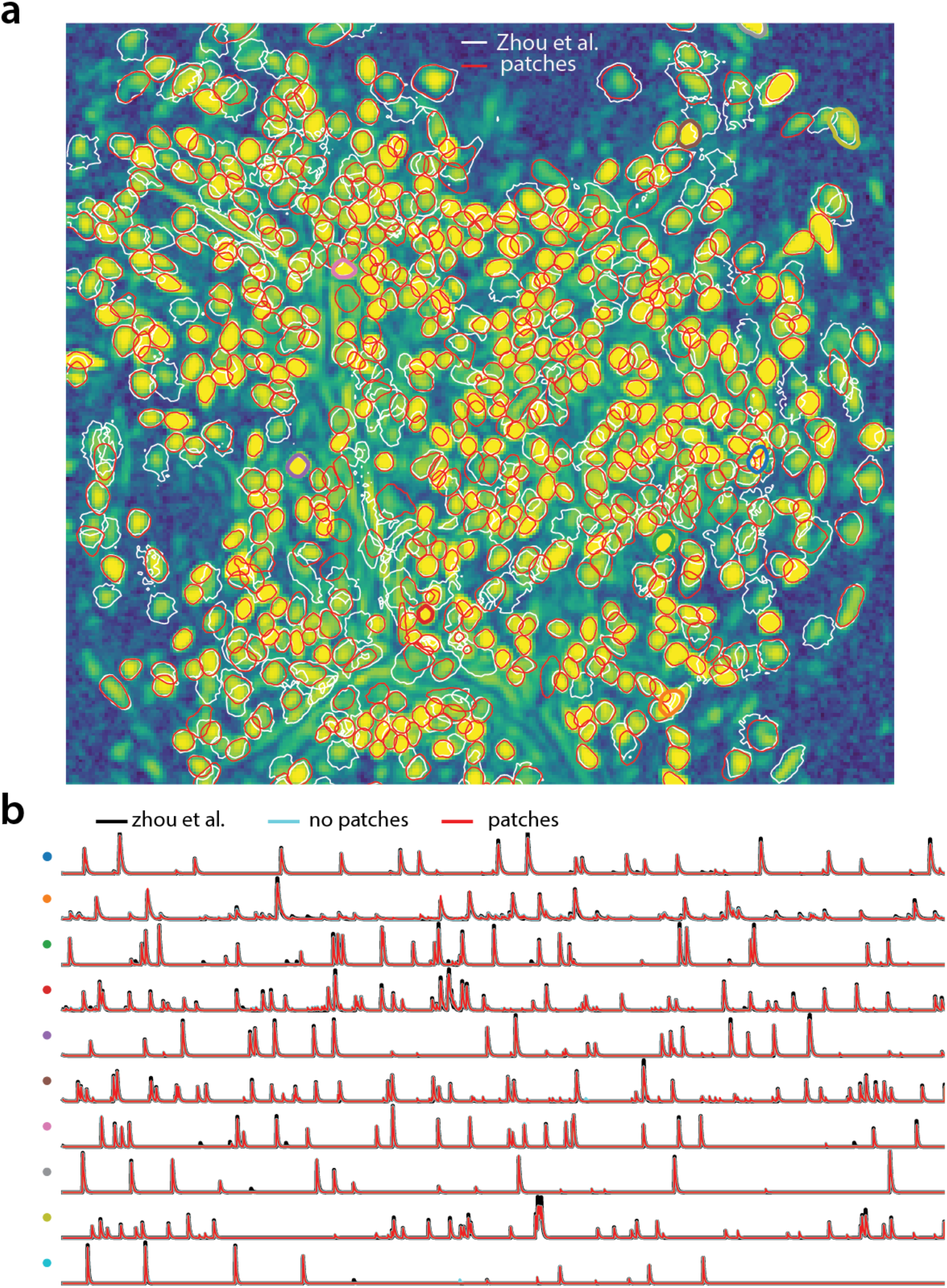
Analyzing microendoscopic 1p data with the CNMF-E algorithm using CaImAn batch. (a) Contour plots of all neurons detected by the CNMF-E (white) implementation of *Zhou et al. (2018)* and CaImAn batch (red) using patches. Colors match the example traces shown in (b), which illustrate the temporal components of 10 example neurons detected by both implementations. CaImAn batch reproduces with reasonable fidelity the results of *Zhou et al. (2018)*.

**Figure 8.**
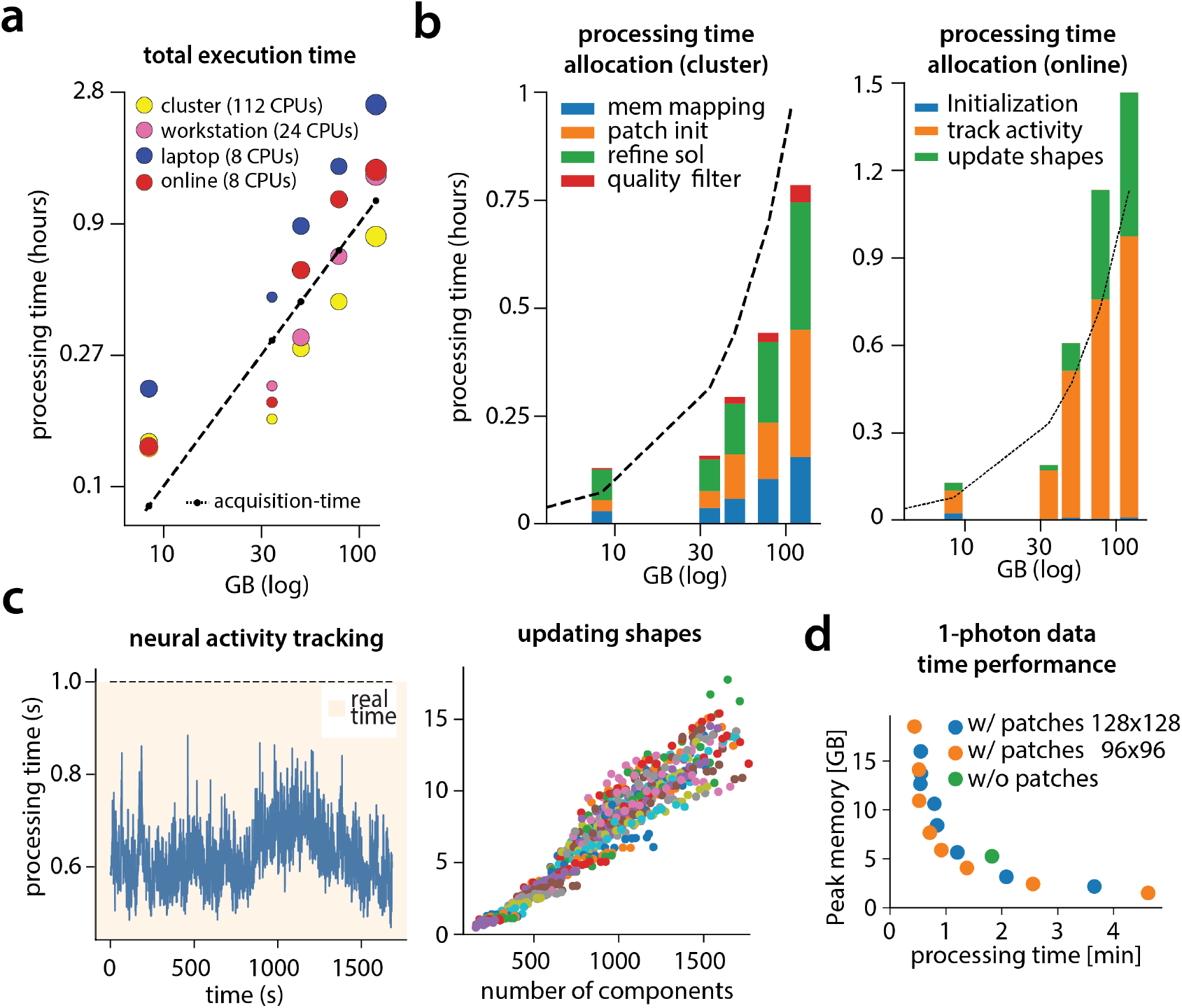
Time performance of CaImAn batch and CaImAn online for the analyzed datasets. (a) Log-log plot of total processing time as a function of data size for CaImAn batch for the 5 largest two-photon datasets using three different processing infrastructures: i) a laptop with 8 CPUs (blue), ii) a desktop workstation with 24 CPUs (magenta), and iii) a HPC where 112 CPUs are allocated (yellow). The results indicate a near linear scaling of the processing time with the size of dataset, with additional dependence on the number of found neurons (size of each point). Even very large datasets (> 100GB) can be processed efficiently with a single laptop, whereas access to a HPC enables processing with speed faster than the acquisition time (considered 30Hz for a 512×512 FOV here). The results of CaImAn online using the laptop are also plotted in red indicating near real-time processing speed. (b) Break down of processing time for CaImAn batch (left) and CaImAn online (right) (excluding motion correction). (Left) Processing with CNMF in patches and refinement takes most of the time for CaImAn batch. Right: Tracking neural activity and new neuron detection can be done in real-time for CaImAn online. (c) (Left) Cost of neural activity online tracking for the whole brain zebrafish dataset (maximum time over all planes per frame). Tracking can be done in real-time. (Right) The most expensive part during online processing occurs while updating the spatial footprints, a step that can be distributed or parallelized. Each color corresponds to the update cost for the various different planes. (d) Cost analysis of CNMF-E implementation for processing a 6000 frames long 1p dataset. Processing in patches in parallel induces a time/memory tradeoff and can lead to speed gains (patch size in legend).

### Analyzing 1p microendoscopic data using CaImAn

We tested the CNMF-E implementation of CaImAn batch on *in vivo* microendosopic data from mouse dorsal striatum, with neurons expressing GCaMP6f. 6000 frames were acquired at 30 frames per second while the mouse was freely moving in an open field arena (for further details refer to *Zhou et al. (2018)*). In Fig. 7 we report the results of the analysis using CaImAn batch with patches and compare to the results of the MATLAB^®^ implementation of *Zhou et al. (2018)*. Both implementations detect similar components (Fig. 7a) with an F_l_-score of 0.89. 573 neurons were found by both implementations. 106 and 31 additional components were detected by *Zhou et al. (2018)* and CaImAn batch respectively. The median correlation between the temporal traces of neurons detected by both implementations was 0.86. Similar results were also obtained by running CaImAn without patches. Ten example temporal traces are plotted in Fig. 7b.

### Computational performance of CaImAn

We examined the performance of CaImAn in terms of processing time for the various analyzed datasets presented above (Fig. 8). The processing time discussed here excludes motion correction, which is highly efficient and primarily depends on the level of the FOV discretization for non-rigid motion correction (*Pnevmatikakis and Giovannucci, 2017*). For CaImAn batch, each dataset was analyzed using three different computing architectures: i) a single laptop (MacBook Pro) with 8 CPUs and 16GB of RAM (blue in Fig. 8a), ii) a linux-based workstation (CentOS) with 24 CPUs and 128GB of RAM (magenta), and iii) a linux-based HPC cluster (CentOS) where 112 CPUs (4 nodes, 28 CPUs each) were allocated for the processing task (yellow). Fig. 8a shows the processing of CaImAn batch as a function of dataset size on the 5 longest datasets, whose size exceeded 8GB, on log-log plot.

Results show that, as expected, employing more processing power results in faster processing. CaImAn batch on a HPC cluster processes data faster than acquisition time (Fig. 8a) even for very large datasets. Processing of an hour long dataset was feasible within 3 hours on a single laptop, even though the dataset has size multiple times the available RAM memory. Here, acquisition time is defined as number of frames times imaging rate, computed based on the assumption of imaging a FOV discretized over a 512 × 512 grid at a 30Hz rate (a typical two-photon imaging setup with resonant scanning microscopes), and a representation of the measurements using single precision arithmetic, which is the minimum precision required for standard algebraic processing. These assumptions lead to a data rate of ~105GB/hour. In general the performance scales linearly with the number of frames (and hence, the size of the dataset), but a dependence is also observed with respect to the number of components. The majority of the time (Fig. 8b-left) the majority of the time required for CaImAn batch processing is taken by CNMF algorithmic processing either during the initialization in patches (orange bar) or during merging and refining the results of the individual patches (green bar).

Fig. 8a also shows the speed performance of CaImAn online (red markers). Because of the low memory requirements of the streaming algorithm, this performance only mildly depends on the computing infrastructure allowing for near real-time processing speeds on a standard laptop (Fig. 8a). As discussed in *Giovannucci et al*. (*2017*) processing time of CaImAn online depends primarily on i) the computational cost of tracking the temporal activity of discovered neurons, ii) the cost of detecting and incorporating new neurons, and iii) the cost of periodic updates of spatial footprints. Fig. 8b-right shows that the two first steps, which are required for each frame, can be done in real-time. In Fig. 8c the cost per frame is plotted for the analysis of the whole brain zebrafish recording. The lower imaging rate (1Hz) allows for the tracking of neural activity to be done with computational cost significantly lower than the 1 second between volume imaging time (Fig. 8c), even in the presence of a large number of components (typically more than 1000 per plane, Fig. 6) and the significantly larger FOV (2048 × 1188 pixels). As expected the cost of updating spatial footprints can be significantly larger if done simultaneously for all components (Fig. 8c, bottom). However, the average cost of updating a single spatial footprint is roughly 8ms, enabling real-time processing *for each* frame, when this step is evenly distributed among different frames/volumes, or is performed by a parallel independent process (*Giovannucci et al., 2017*).

The cost of processing 1p data in CaImAn batch using the CNMF-E algorithm (*Zhou et al., 2018*) is shown (Fig. 8d) for the workstation hardware. Splitting in patches and processing in parallel can lead to computational gains at the expense of increased memory usage. This is because the CNMF-E introduces a background term that has the size of the dataset and needs to be loaded and updated in memory in two copies. This leads to processing times that are slower compared to the standard processing of 2p datasets, and higher memory requirements. However, as Fig. 8d demonstrates, memory usage can be controlled enabling scalable inference at the expense of slower processing speeds.

### CaImAn successfully tracks neurons across multiple days

Fig. 9 shows an example of tracking neurons across 6 different sessions corresponding to 6 different days of mouse cortex *in vivo* data using our multi-day registration algorithm RegisterMulti (see Methods, Alg. 6). 453, 393, 375, 378, 376, and 373 active components were found in the six sessions, respectively. Our tracking method detected a total of 686 distinct active components. Of these, 172, 108, 70, 92, 82, and 162 appeared in exactly 1, 2, 3, 4, 5, and all 6 sessions respectively. Contour plots of the 162 components that appeared in all sessions are shown in Fig. 9a, and parts of the FOV are highlighted in Fig. 9d showing that components can be tracked in the presence of non-rigid deformations of the FOV between the different sessions.

**Figure 9.**
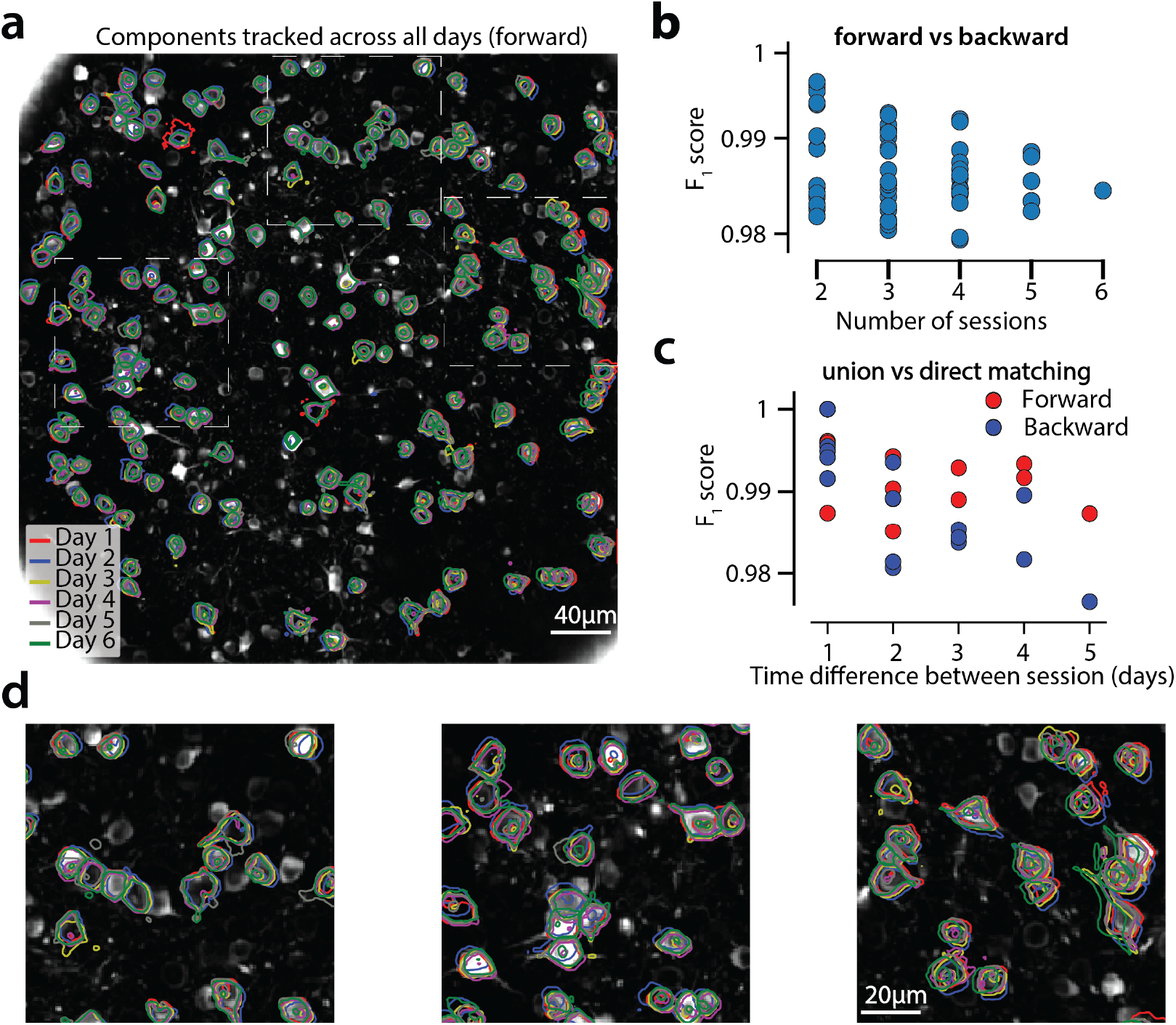
Components registered across six different sessions (days). (a) Contour plots of neurons that were detected to be active in all six imaging sessions overlaid on the correlation image of the sixth imaging session. Each color corresponds to a different session. (b) Stability of multiday registration method. Comparisons of forward and backward registrations in terms of *F*_1_ scores for all possible subsets of sessions. The comparisons agree to a very high level indicating the stability of the proposed approach. (c) Comparison (in terms of *F*_1_ score) of pair-wise alignments using readouts from the union vs direct alignment. The comparison is performed for both the forward and the backwards alignment. For all pairs of sessions the alignment using the proposed method gives very similar results compared to direct pairwise alignment. (d) Magnified version of the tracked neurons corresponding to the squares marked in panel (a). Neurons in different parts of the FOV exhibit different shift patterns over of the course of multiple days, but can nevertheless be tracked accurately by the proposed multiday registration method.

To test the stability of RegisterMulti for each subset of sessions, we repeated the same procedure running backwards in time starting from day 6 and ending at day 1, a process that also generated a total of 686 distinct active components. We identified the components present in *at least* a given subset of sessions when using the forward pass, and separately when using the backwards pass, and compared them against each other (Fig. 9b) for all possible subsets. Results indicate a very high level of agreement between the two approaches with many of the disagreements arising near the boundaries (data not shown). Disagreements near the boundaries can arise because the forward pass aligns the union with the FOV of the last session, whereas the backwards pass with the FOV of the first session, potentially leading to loss of information near the boundaries.

A step by step demonstration of the tracking algorithm for the first three sessions is shown in the appendix (Fig. 10). Our approach allows for the comparison of two non-consecutive sessions through the union of components without the need of a direct pairwise registration (Fig. 10f), where it is shown that registering sessions 1 and 3 directly and through the union leads to nearly identical results. Fig. 9c compares the registrations for all pairs of sessions using the forward (red) or the backward (blue) approach, with the direct pairwise registrations. Again, the results indicate a very high level of agreement, indicating the stability and effectiveness of the proposed approach.

**Appendix 0 Figure 10.**
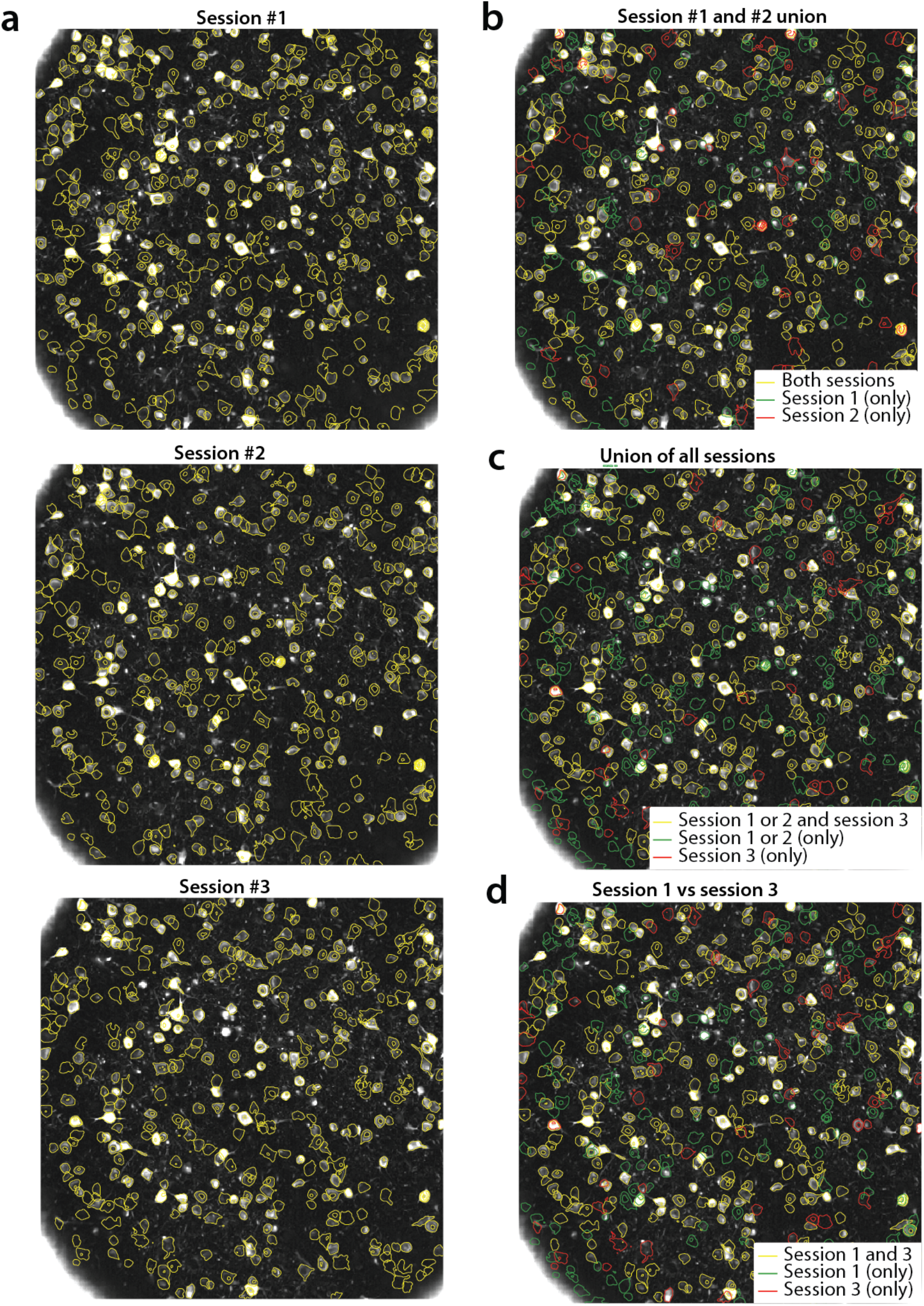
Tracking neurons across days, step-by-step description of multi session registration (Fig. 9). (a) Correlation image overlaid to contour plots of the neurons identified by CaImAn batch in day 1 (top, 453 neurons), 2 (middle, 393 neurons) and 3 (bottom, 375 neurons). (b) Result of the pairwise registration between session 1 and 2. The union of distinct active components consists of the components that were active in i) both sessions (yellow - where only the components of session 2 are displayed), ii) only in session 2 (green), and iii) only in session 1, aligned to the FOV of session 2 (red). (c) At the next step the union of sessions 1 and 2 is registered with the results of session 3 to produce the union of all distinct components aligned to the FOV of session 3. (d) Comparison of non-consecutive sessions without pairwise registration. Keeping track of which session each component was active in, enables efficient and stable comparisons.

**Appendix 0 Figure 11.**
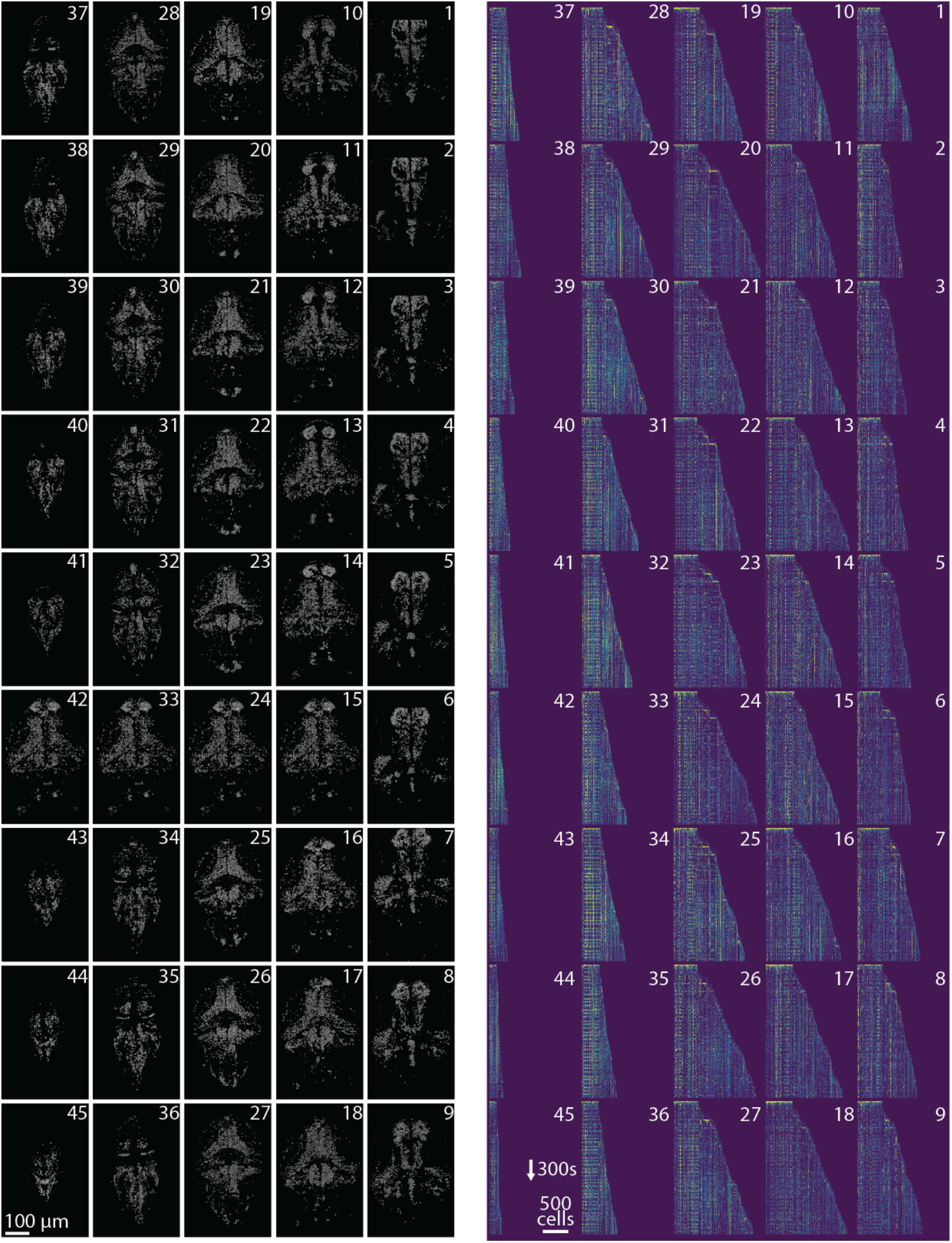
Profile of spatial (left) and temporal (right) components found in each plane of the whole brain zebrafish recording. (Left) Components are extracted with CaImAn online and then max-thresholded. (Right) See Results section for a complete discussion.

## Discussion

### Reproducible and scalable analysis for the 99%

Significant advances in the reporting fidelity of fluorescent indicators, and the ability to simultaneously record and modulate neurons granted by progress in optical technology, have propelled calcium imaging to being the main experimental method in systems neuroscience alongside electrophysiology recordings. The resulting increased adoption rate has generated an unprecedented wealth of imaging data which poses significant analysis challenges. The goal of CaImAn is to provide the experimentalist with a complete suite of tools for analyzing this data in a formal, scalable, and reproducible way. The goal of this paper is to present the features of CaImAn and examine its performance in detail. CaImAn embeds existing methods for preprocessing calcium imaging data into a MapReduce framework and augments them with supervised learning algorithms and validation metrics. It builds on the CNMF algorithm of *Pnevmatikakis et al*. (*2016*) for source extraction and deconvolution, extending it along the lines of i) reproducibility and performance improvement, by automating quality assessment through the use of unsupervised and supervised learning algorithms for component detection and classification, and ii) scalability, by enabling fast large scale processing with standard computing infrastructure (e.g., a commodity laptop or workstation). Scalability is achieved by either using a MapReduce batch approach, which employs parallel processing of spatially overlapping, memory mapped, data patches; or by integrating the online processing framework of *Giovannucci et al*. (*2017*) within our pipeline. Apart from computational gains both approaches also result in improved performance. Towards our goal of providing a single package for dealing with standard problems arising in analysis of imaging data, CaImAn also includes an implementation of the CNMF-E algorithm of *Zhou et al. (2018)* for the analysis of microendoscopic data, as well as with a novel method for registering analysis results across multiple days.

### Towards surpassing human neuron detection performance

To evaluate the performance of CaImAn batch and CaImAn online, we generated a corpus of multiply annotated two-photon imaging datasets. The results indicated a surprising level of disagreement between individual labelers, highlighting both the difficulty of the problem, and the non-reproducibility of the laborious task of human annotation. CaImAn reached near-human performance with respect to this ground truth, by using the *same* parameters for all the datasets without dataset dependent parameter tweaking. Such tweaking could for example include setting the SNR threshold based on the noise level of the recording, the complexity of the neuropil signal based on the level of background activity, or specialized treatment around the boundaries of the FOV to compensate for eventual imaging artifacts.

Apart from being used as a benchmarking tool, the set of manual annotations can also be used as labeled data for supervised learning algorithms. CaImAn uses two CNN based classifiers trained on (a subset of) this data, one for post processing component classification in CaImAn batch, and the other for detecting new neurons in residual images in the CaImAn online. The deployment of these classifiers resulted in significant gains in terms of performance, and we expect further advances in the future. The annotations will be made freely available to the community upon publication of the paper for benchmarking and training purposes.

### CaImAn batch vs CaImAn online

Our results suggest similar performance between CaImAn batch and CaImAn onlinein terms of processing speed and quality of results with CaImAn online outperforming CaImAn batch on longer datasets in terms of neuron detection, possibly due to its inherent ability to adapt to non-stationarities arising during the course of a large experiment, and underperforming on shorter datasets potentially due to lack of enough information. By contrast, CaImAn batch extracts better traces compared to CaImAn online with respect to “ground truth” traces. While multiple passes over the data with CaImAn online can mitigate these shortcomings, this still depends on good initialization with CaImAn batch, as the analysis of the whole brain zebrafish dataset indicates. In offline setups, CaImAn online could also benefit from the post processing component evaluation tools used in batch mode. e.g., using the batch classifier for detecting false positive components at the end of the experiment.

What sets the two algorithms apart is the streaming processing mode of CaImAn online which, besides lowering memory requirements, can be used to enable novel types of closed-loop all-optical experiments (*Packer et al., 2015; Carrillo-Reid et al., 2017*). As discussed in *Giovannucci et al*. (*2017*), typical all-optical closed-loop experiments require the pre-determination of ROIs that are monitored/modulated. Processing with CaImAn online can improve upon this by allowing identification and modulation of new neurons on the fly, greatly expanding the space of possible experiments. Even though our simulated online processing setup is not integrated with hardware to an optical experimental setup, our results indicate that CaImAn online performed close to real-time in most cases, without optimizing for speed. This suggest that large scale closed-loop experiments with single cell resolution are feasible by combining existing all-optical technology and our proposed analysis method.

### Future directions

While CaImAn uses a highly scalable processing pipeline for two-photon datasets, processing of one-photon microendoscopic imaging data is less scalable due to the more complex background model that needs to be retained in memory during processing. Adapting CaImAn online to the one-photon data processing algorithm of *Zhou et al. (2018)* is a promising way for scaling up efficient processing in this case. The continuing development and quality improvement of neural activity indicators has enabled direct imaging of neural processes (axons/dendrites), imaging of synaptic activity (*Xie et al., 2016*), or direct imaging of voltage activity *in vivo* conditions (*Piatkevich et al., 2018*). While the approach presented here is tuned for somatic imaging through the use of various assumptions (space localized activity, CNN classifiers trained on images of somatic activity), the technology of CaImAn is largely transferable to these domains as well. These extensions will be pursued in future work.

## Methods and Materials

### Memory mapping

In order to efficiently access data in parallel, CaImAn batch relies on memory mapping. With memory mapped (mmap) arrays, arithmetic operations can be performed on data residing on the hard drive without explicitly loading it to RAM, and slices of data can be indexed and accessed without loading the full file in memory, enabling out-of-core processing (*Toledo, 1999*). The order in which data in a memory mapped file is stored on the hard drive can dramatically affect the read-write performance of out-of-core operations on spinning disks, and to a lesser degree on solid state drives. On modern computers tensors are stored in linear format, no matter the number of the array dimensions. Therefore, one has to decide which elements of an array are contiguous in memory: in *row-major order*, consecutive elements of a row (first-dimension) are next to each other, whereas in *column-major order* consecutive elements of a column (last dimension) are contiguous. Such decisions significantly affect the speed at which data is read or written: in *column-major order* reading a full column is fast because memory is read in a single sequential block, whereas reading a row is inefficient since only one element can be read at a time and all the data needs to be accessed. Therefore, the original dataset must be saved in the right order to avoid performance problems.

In the context of calcium imaging datasets, CaImAn batch represents the datasets in a matrix form *Y*, where each row corresponds to a different imaged pixel, and each column to a different frame. As a result, a *column-major order* mmap file enables the fast access of individual frames at a given time, whereas a *row-major order* files enables the fast access of an individual pixel at all times. To facilitate processing in patches CaImAn batch stores the data in *row-major order*. In practice, this is opposite to the order with which the data appears, one frame at a time. In order to reduce memory usage and speed up computation CaImAn batch employs a MapReduce approach, where either multiple files or multiple chunks of a big file composing the original datasets are processed and saved in mmap format in parallel. This operation includes two phases, first the chunks/files are saved in multiple row-major mmap format, and then chunks are simultaneously combined into a single large row-major mmap file. In order to reduce preprocessing steps, if the file(s) need to be corrected for motion artifacts, chunks of the registered data can be stored on-the-fly during motion correction.

### Mathematical model of the CNMF framework

The CNMF framework (Fig. 1d) for calcium imaging data representation can be expressed in mathematical terms as (*Pnevmatikakis et al., 2016*)

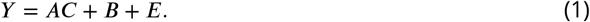

Here, 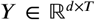 denotes the observed data written in matrix form, where *d* is the total number of observed pixels/voxels, and *T* is the total number of observed timesteps (frames). 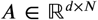 denotes the matrix of the *N* spatial footprints, *A* = [**a**_1_, **a**_2_, …, **a**_*N*_], with 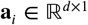 being the spatial footprint of component *i*. 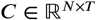 denotes the matrix of temporal components, *C* = [**c**_1_, **c**_2_, …, **c**_*n*_]^Τ^, with 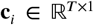 being the temporal trace of component *i*. ***B*** is the background/neuropil activity matrix. For two-photon data it is modeled as a low rank matrix *B* = **bf**, where 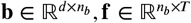 correspond to the matrices of spatial and temporal background components, and *n_b_* is the number of background components. For the case of micro-endoscopic data the integration volume is much larger and the low rank model is inadequate. A solution comes from the CNMF-E algorithm of *Zhou et al*. (*2018*) where the background is modeled as

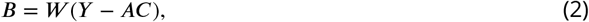

where 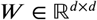 is an appropriate weight matrix, where the (*i*, *j*) entry models the influence of the neuropil signal of pixel *j* to the neuropil signal at pixel *i*.

### Combining results from different patches

To combine results from the different patches we first need to account for the overlap at the boundaries. Neurons lying close to the boundary between neighboring patches can appear multiple times and must be merged. With this goal, we optimized the merging approach used in *Pnevmatikakis et al*. (*2016*): Groups of components with spatially overlapping footprints whose temporal traces are correlated above a threshold are replaced with a single component, that tries to explain as much of the variance already explained by the “local” components (as opposed to the variance of the data as performed in (*Pnevmatikakis et al., 2016*)). If *A*_old_, *C*_old_ are the matrices of components to be merged, then the merged component **a**_*m*_, **c**_*m*_ are given by the solution of the rank-1 NMF problem:

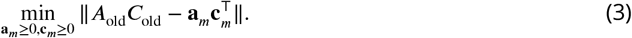

Prior to merging, the value of each component at each pixel is normalized by the number of patches that overlap in this pixel, to avoid counting the activity of each pixel multiple times.

We follow a similar procedure for the background/neuropil signals from the different patches. For the case of two-photon data, the spatial background/neuropil components for each patch can be updated by keeping their spatial extent intact to retain a local neuropil structure, or they can be merged when they are sufficiently correlated in time as described above to promote a more global structure. For the case of one-photon data, CNMF-E estimates the background using a local auto-regressive process (see Eq. (2)) *Zhou et al., 2018*), a setup that cannot be immediately propagated when combining the different patches. To combine backgrounds from the different patches, we first approximate the backgrounds ***B**^i^* from all the patches *i* with a low rank matrix using non-negative matrix factorization of rank *g_b_* to obtain global spatial, and temporal background components.

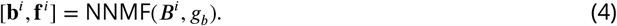

The resulting components are embedded into a large matrix 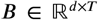 that retains a low rank structure. After the components and backgrounds from all the patches have been combined, they are further refined by running CNMF iteration of updating spatial footprints, temporal traces, and neuropil activity. CaImAn batch implements these steps in a highly parallel fashion (as also described in *Pnevmatikakis et al*. (*2016*)): Temporal traces whose corresponding spatial traces do not overlap can be updated in parallel. Similarly, the rows of the matrix of spatial footprints ***A*** can also be updated in parallel (2b). The process is summarized in algorithmic format in Alg. 1.

### Initialization strategies

Source extraction using matrix factorization requires solving a bi-convex problem where initialization plays a critical role. The CNMF/CNMF-E algorithms use initialization methods that exploit the locality of the spatial footprints to efficiently identify the locations of candidate components (*Pnevmatikakis et al., 2016*; *Zhou et al., 2018*). CaImAn incorporates these methods, extending them by using the temporal locality of the calcium transient events. The available initialization methods for CaImAn batch include:

**GreedyROI:** This approach, introduced in *Pnevmatikakis et al. (2016)*, first spatially smooths the data with a Gaussian kernel of size comparable to the average neuron radius, and then initializes candidate components around locations where maximum variance (of the smoothed data) is explained. This initialization strategy is fast but requires specification of the number of components by the user.
**RollingGreedyROI:** The approach, introduced in this paper, operates like GreedyROI by spatially smoothing the data and looking for points of maximum variance. Instead of working across all the data, RollingGreedyROI looks for points of maximum variance on a rolling window of a fixed duration, e.g., 3 seconds, and initializes components by performing a rank one NMF on a local spatial neighborhood. By focusing into smaller rolling windows, RollingGreedyROI can better isolate single transient events, and as a result detect better neurons with sparse activity. RollingGreedyROI is the default choice for processing of 2-photon data.
**GreedyCorr:** This approach, introduced in *Zhou et al. (2018)*, initializes candidate components around locations that correspond to the local maxima of an image formed by the pointwise product between the correlation image and the peak signal-to-noise ratio image. By setting a threshold for acceptance, this approach does not require the prior specification of number of components. This comes at the expense of a higher computational cost. GreedyCorr is the default choice for processing of 1-photon data.
**SparseNMF:** Sparse NMF approaches, when ran in small patches, can be effective for quickly uncovering spatial structure in the imaging data, especially for neural processes (axons/dendrites) whose shape cannot be easily parametrized and/or localized.

### Algorithm seeding with binary masks

Often locations of components are known either from manual annotation or from labeled data obtained in a different way, such as data from a static structural channel recorded concurrently with the functional indicator. CaImAn can be seeded with binary (or real valued) masks for the spatial footprints. Apart from A, these masks can be used to initialize all the other relevant matrices *C* and *B* as well. This is performed by i) first estimating the temporal background components **f** using only data from parts of the FOV not covered by any masks and, ii) then estimating the spatial background components **b**, and then estimating *A, C* (with *A* restricted to be non-zero only at the locations of the binary masks), using a simple NMF approach. Details are given in Alg. 2.

### Details of quality assessment tests

Here we present the unsupervised and supervised quality assessment tests in more detail (Fig. 2).

#### Matching spatial footprints to the raw data

Let **a**_*i*_., **c**_*i*_ denote the spatial footprint and temporal trace of component *i*, and the let *A_\i_*, *C_\i_* denote the matrices *A, C* when the component *i* has been removed. Similarly, let *Y_i_* = *Y* − *A_\i_*, *C_\i_* − ***B*** denote the entire dataset when the background and the contribution of all components except *i* have been removed. If component *i* is real then *Y_i_* and 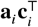 will look similar during the time intervals when the component *i* is active. As a first test CaImAn finds the first *N_p_* local peaks of *c_i_* (e.g., *N_p_* = 5), constructs intervals around these peaks, (e.g., 50 ms in the past and 300ms in the future, to cover the main part of a possible calcium transient around that point), and then averages *Y_i_* across time over the union of these intervals to obtain a spatial image < *Y_i_* > (Fig. c). The Pearson’s correlation over space between < *Y_i_* > and **a***_i_*, (both restricted on a small neighborhood around the centroid of **a***_i_*) is then computed, and component *i* is rejected if the correlation coefficient is below a threshold value *θ*_sp_, (e.g., *θ*_sp_ < 0.5). Note that a similar test is used in the online approach of *Giovannucci et al*. (*2017*) to accept for possible new components.

#### Detecting fluorescence traces with high SNR

For a candidate component to correspond to an active neuron its trace must exhibit dynamics reminiscent of the calcium indicator’s transient. A criterion for this can be obtained by requiring the average SNR of trace **c***_i_* over the course a transient to be above a certain threshold *θ*_SNR_, e.g., *θ*_SNR_ = 2, (Fig. 2d). The average SNR is as a measure of how unlikely it is for the transients of **c***_i_* (after some appropriate z-scoring) to have been a result of a white noise process.

To compute the SNR of a trace, let *R = Y − AC − B* be the residual spatiotemporal signal. We can obtain the residual signal for each component *i*, **r***_i_*, by projecting *R* into the spatial footprint **a**^*i*^:

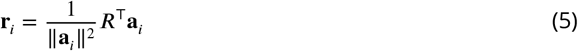

Then the trace **c***_i_* + **r***_i_* corresponds to the non-denoised trace of component *i*. To calculate its SNR we first compute a type of z-score:

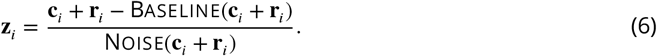

The Baseline(·) function determines the baseline of the trace, which can be varying in the case of long datasets exhibiting baseline trends, e.g., due to bleaching. The function Noise(·) estimates the noise level of the trace. Since calcium transients around the baseline can only be positive, we estimate the noise level by restricting our attention only to the points *t_n_* where c_*i*_ + **r***_i_* is below the baseline value, i.e., *t_n_* = {*t*: **c***_i_*(*t*) + **r***_i_*(*t*) ≤ Baseline(**c***_i_* + **r***_i_*)}, and compute the noise level as the scale parameter of a half-normal distribution (Fig. 2b):

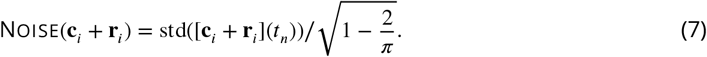

We then determine how likely is that the positive excursions of **z**_*i*_ can be attributed just to noise. We compute the probabilities **p***_i_*(*t*) = Φ(−**z**_*i*_(*t*)), where Φ(·) denotes the cumulative distribution function of a standard normal distribution, and compute the most unlikely excursion over a window of *N_s_* timesteps that corresponds to the length of a typical transient, e.g., *N_s_* = [0.4*s* × *F*], where 0.4s could correspond to the typical length of a GCaMP6f transient, and *F* is the imaging rate.

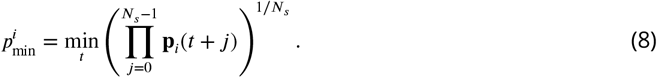

The (averaged peak) SNR of component *i* can then be defined as

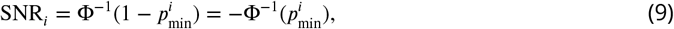

where Φ^-1^ is the quantile function for the standard normal distribution (logit function) and a component is accepted if SNR*_i_* ≥ *θ*_SNR_. Note that for numerical stability we compute 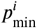 in the logarithmic domain and check the condition 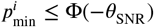.

We can also use a similar test for the significance of the time traces in the spike domain after performing deconvolution. In this case, traces can be considered as spiking if the maximum height due to a spike transient is significantly larger than a threshold. If we assume that the shape of each calcium transient has been normalized to have maximum amplitude 1, then this corresponds to testing ||**s**_*i*_||_∞_ > *θ*_SNR_*σ_i_*, where **s***_i_* represents the deconvolved activity trace for component *i*, and *θ*_SNR_ is again an appropriate SNR threshold, e.g., *θ*_SNR_ = 2, and *σ_i_* is the noise level for trace *i*.

#### Classification through convolutional neural networks (CNNs)

The tests described above are unsupervised but require fine-tuning of two threshold parameters (*θ_sp_*, *θ*_SNR_) that might be dataset dependent and might be sensitive to strong non-stationarities. As a third test we trained a 4-layer CNN to classify the spatial footprints into true or false components, where a true component here corresponds to a spatial footprint that resembles the soma of a neuron (See Fig. 2e and section *Classification through convolutional networks* for details). A simple threshold *θ*_CNN_ can be used to tune the classifier (e.g., *θ*_CNN_ = 0.5).

### Collection of manual annotations and ground truth

We collected manual annotations from four independent labelers who were instructed to find round or donut shaped neurons of similar size using the ImageJ Cell Magic Wand tool *Walker* (*2014*). We focused on manually annotating only cells that were active within each dataset and for that reason the labelers were provided with two summary statistics: i) A movie obtained by removing a running 20th percentile (as a crude background approximation) and downsampling in time by a factor of 10, and ii) the max-correlation image. The correlation image (CI) at every pixel is equal to the average temporal correlation coefficient between that pixel and its neighbors *Smith and Häusser* (*2010*) (8 neighbors were used for our analysis). The max-correlation image is obtained by computing the CI for each batch of 33 seconds (1000 frames for a 30Hz acquisition rate), and then taking the maximum over all these images. Neurons that are inactive during the course of the dataset will be suppressed both from the baseline removed video (since their activity will always be around their baseline), and from the max-correlation image since the variation around this baseline will mostly be due to noise leading to practically uncorrelated neighboring pixels. 9 different mouse in vivo datasets were used from various brain areas and labs. A description is given in Table 2. To create the consensus ground truth, the labelers were asked to jointly resolve the inconsistencies with each others annotations.

The annotation procedure provides a binary mask per selected component. On the other hand, the output of CaImAn for each component is a non-negatively valued vector over the FOV (a real-valued mask). The two sets of masks differ not only in their variable type but also in their general shape: Manual annotation through the Cell Magic Wand tool tends to produce circular shapes, whereas the output of CaImAn will try to accurately estimate the shape of each active component. To construct ground truth that can be directly used for comparison, the binary masks from the manual annotations were used to seed the CNMF algorithm (Alg. 2). This produced a set of ground truth real valued components with spatial footprints restricted to the areas provided by the annotations, and a corresponding set of temporal components that can be used to evaluate the performance of CaImAn (Fig. 4). Registration was performed using the RegisterPair algorithm (Alg. 5) and match was counted as a true positive when the (modified)Jaccard distance (Eq. 11) was below 0.7. Details of the registration procedure are given below (see *Component registration*).

### Classification through convolutional neural networks (CNNs)

CaImAn uses two CNN classifiers; one for post processing component screening in CaImAn batch, and a different one for screening candidate components in CaImAn online. In both cases a 4 layer CNN was used, with architecture as described in Fig. 2e.

#### CaImAn batch classifier for post processing classification

The purpose of the batch classifier is to classify the components detected by CaImAn batch into neuron somas or other shapes, by examining their spatial footprints. Only three annotated datasets (.03.00.t, NF.04.00.t, NF.02.00) were used to train the batch classifier. The set of estimated footprints from running CaImAn batch initialized with the consensus ground truth was matched to the set of ground truth footprints. Footprints matched to ground truth components were considered positive examples, whereas the remaining components were labeled as negatives. The two sets were enriched using data augmentation (rotations, reflections, contrast manipulation etc.) through the Keras library (keras.io) and the CNN was trained on 60% of the data, leaving 20% for validation and 20% for testing. The CNN classifier reached an accuracy of 97% on test data; that also generalized to the rest of the datasets (Fig. 2e) without any parameter change.

#### Online classifier for new component detection

The purpose of the CaImAn online classifier is to detect new components based on their spatial footprints by looking at the mean across time of the residual buffer. To construct the ground truth data for the online classifier, CaImAn batch was run on the first five annotated datasets seeded with the masks obtained through the manual annotations. Subsequently the activity of random subsets of found components and the background was removed from contiguous frames of the raw datasets to construct residual buffers, which were averaged across time. From the resulting images patches were extracted corresponding to positive examples (patches around a neuron that was active during the buffer) and negative examples (patches around other positions within the FOV). A neuron was considered active if its trace attained an average peak-SNR value of 4 or higher during the buffer interval. Similarly to the batch classifier, the two sets were augmented and split into training, validation and testing sets. The resulting classifier reached a 98% accuracy on the testing set, and also generalized well when applied to different datasets.

#### Differences between the two classifiers

Although both classifiers examine the spatial footprints of candidate components, their required performance characteristics are different which led us to train them separately. The batch classifier examines each component as a post-processing step to determine whether its shape corresponds to a neural cell body. As such, false positive and false negative examples are treated equally and possible mis-classifications do not directly affect the traces of the other components. By contrast, the online classifier operates as part of the online processing pipeline. In this case, a new component that is not detected in a residual buffer is likely to be detected later should it become more active. On the other hand, a component that is falsely detected and incorporated in the online processing pipeline will continue to affect the future buffer residuals and the detection of future components. As such the online algorithm is more sensitive to false positives than false negatives. To ensure a small number of false positive examples under testing conditions, only components with average peak-SNR value at least 4 were considered as positive examples during training of the online classifier.

### Component registration

Fluorescence microscopy methods enable imaging the same part of the brain across different sessions that can span multiple days or weeks. While the microscope can visit the same location in the brain with reasonably high precision, the FOV might might not precisely match due to misalignments or deformations in the brain medium. CaImAn provides routines for FOV alignment and component registration across multiple sessions/days. Let 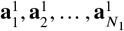 and 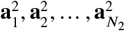 the sets of spatial components from sessions 1 and 2 respectively, where *N*_1_ and *N*_2_ denote the total number of components from each session. We first compute the FOV displacement by aligning some summary images from the two sessions (e.g., mean or correlation image), using some non-rigid registration method, e.g., NoRMCorre (*Pnevmatikakis and Giovannucci, 2017*). We apply the estimated displacement field to the components of *A*_1_ to align them with the FOV of session 2. To perform the registration, we construct a pairwise distance matrix 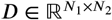 with 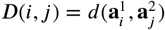, where *d*(·, ·) denotes a distance metric between two components. The chosen distance corresponds to theJaccard distance between the binarized versions of the components. A real valued component **a** is converted into its binary version *m*(**x**) by setting to 1 only the values of **a** that are above the maximum value of a times a threshold *θ_b_*, e.g., *θ_b_* = 0.2:

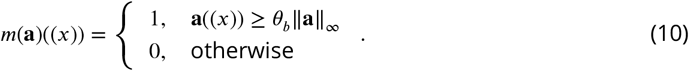

To compute the distance between two binary masks *m*_1_, *m*_2_, we use theJaccard index (intersection over union) which is defined as

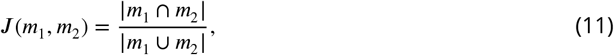

and use it to define the distance metric as

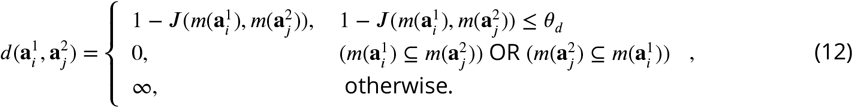

where *θ_d_* is a distance threshold, e.g., 0.5 above which two components are considered nonmatching and their distance is set to infinity to prevent false assignments.

After the distance matrix *D* has been completed, an optimal matching between the components of the two sessions is computed using the Hungarian algorithm to solve the linear assignment problem. As infinite distances are allowed, it is possible to have components from both sessions that are not matched with any other component. This process of registering components across two sessions (RegisterPair) is summarized in Alg. 5.

To register components across multiple sessions, we first order the sessions chronologically and register session 1 against session 2. From this registration we construct the union of the distinct components between the two sessions by keeping the matched components from session 2 as well as the non-matched components from both sessions aligned to the FOV of session 2. We then register this union of components to the components of session 3 and repeat the procedure until all sessions are have been registered. This process of multi session registration (RegisterMulti) is summarized in Alg. 6. At the end of the process the algorithm produces a list of matches between the components of each session and the union of all active distinct components, allowing for efficient tracking of components across multiple days (Fig. 9), and the comparison of non-consecutive sessions through the union without the need of direct pairwise registration (Fig. 10)). An alternative approach to the problem of multiple session registration (CellReg) was presented recently by *Sheintuch et al*. (*2017*) where the authors register neurons across multiple days by first constructing a similar union set of all the components which is then refined using a clustering procedure. RegisterMulti differs from the CellReg method of *Sheintuch et al. (2017)* in a few key ways, that highlight its simplicity and robustness:

- RegisterMulti uses a very simple intersection over union metric to estimate the distance between two neighboring neurons after the FOV alignment. Cells that have a distance above a given threshold are considered different by default and are not tested for matching. This parameter is intuitive to set a priori for each dataset. In contrast CellReg uses a probabilistic framework based on the joint probability distribution between the distance of two cells and the correlation of their shapes that makes specific parametric assumptions about the distributions of centroid distances between the same and different cells, as well as their shape correlations. This model needs to be re-evaluated for every different set of sessions to be registered and potentially requires a lot of data to learn the appropriate distance metric.
- RegisterMulti uses the Hungarian algorithm to register two different set of components, a practice that solves the linear assignment problem optimally under the assumed distance function. In contrast CellReg uses a greedy method for initializing the assignment of cells to the union superset relying on the following clustering step to refine these estimates, and thus adding extra computational burden to the registration procedure.

### Implementation details for CaImAn batch

Each dataset was processed using the same set of parameters, excepting the expected size of neurons (estimated by inspecting the correlation image), the size of patches and expected number of neurons per patch (estimated by inspecting the correlation image). For the dataset `N.01.01`, where optical modulation was induced, the threshold for merging neurons was slightly higher (the stimulation caused clustered synchronous activity). For shorter datasets, rigid motion correction was sufficient; for longer datasets `K53,J115` we applied non-rigid motion correction. The parameters for the automatic selection of components were optimized using only the first three datasets and fixed for all the remaining files. For all datasets the background neuropil activity was modeled as a rank 2 matrix, and calcium dynamics were modeled as a first order autoregressive process. The remaining parameters were optimized so that all the datasets could be run on a machine with less than 128GB RAM.

### Implementation details for CaImAn online

Datasets were processed for two epochs with the exception of the longer datasets `K53,J115,J123` where only one pass of the data was performed to limit computational cost. For each dataset the online CNN classifier was used to detect new neurons, and five candidate components were considered for each frame. The online CNN classifier had the same threshold 0.5 for all datasets, with the exception of the longest datasets `J115,J123` where the threshold was set to 0.75. Setting the threshold to 0.5 for these datasets led to slightly poorer performance. Large datasets were spatially decimated by a factor of 2 to enhance processing speed, a step that did not lead to changes in detection performance. For all datasets the background neuropil activity was modeled as a rank 2 matrix, and calcium dynamics were modeled as a first order autoregressive process. For each dataset, CaImAn online was initialized on the first 200 frames, using the BareInitialization on the entire FOV with only 2 neurons, so in practice all the neurons were detected during the online mode. To highlight the *truly* online processing mode, no post-processing of the results was used, a step that can further enhance the performance of the algorithm. Similarly to batch processing, the expected size of neurons was chosen separately for each dataset after inspecting the correlation image.

For the analysis of the whole brain zebrafish dataset, CaImAn online was run for 1 epoch with the same parameters as above, with only differences appearing in the number of neurons during initialization (600 vs 2), and the value of the threshold for the online CNN classifier (0.75 vs 0.5). The former decision was motivated by the goal of retrieving with a single pass neurons from a preparation with a denser level of activity over a larger FOV in this short dataset (1885 frames). To this end, the number of candidate neurons at each timestep was set to 10 (per plane). The threshold choice was motivated by the fact that the classifier was trained on mouse data only, and thus a higher threshold choice would help diminish potential false positive components. Rigid motion correction was applied online to each plane.

### Performance quantification as a function of SNR

To quantify performance as a function of SNR we approximate the ground truth traces by running CaImAn batch on the datasets seeded with the “consensus” binary masks obtained from the manual annotators. After that the average peak-SNR of a trace **c** with corresponding residual signal **r** (5) is obtained as

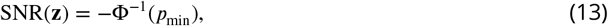

where Φ^−1^(·) denotes the probit function (quantile function for the standard Gaussian distribution), **z** is the z-scored version of **c + r** (6) and *p*_min_ is given by (8).

Let 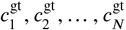 be the ground truth traces and 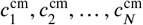 be their corresponding CaImAn inferred traces. Here we assume that false positive and false negative components are matched with trivial components that have 0 SNR. Let also 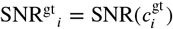 and 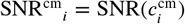, respectively. After we compute the SNR for both ground truth and inferred traces the performance algorithm can be quantified in multiple ways as a function of a SNR thresholds *θ*_SNR_:

**Precision:** Precision at level *θ*_SNR_, can be computed as the fraction of detected components with SNR^cm^ > *θ*_SNR_ that are matched with ground truth components. It quantifies the certainty that a component detected with a given SNR or above corresponds to a true component.

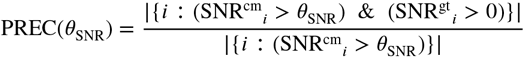
**Recall:** Recall at level *θ*_SNR_, can be computed as the fraction of ground truth components with SNR^gt^ > *θ*_SNR_ that are detected by the algorithm. It quantifies the certainty that a ground truth component with a given SNR or above is detected.

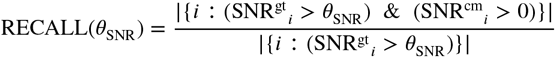
*F*_1_ **score:** An overall *F*_1_ score at level *θ*_SNR_, can be obtained by computing the harmonic mean between precision and recall

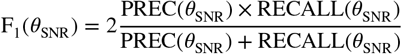 The cautious reader will observe that the precision and recall quantities described above are not computed in the same set of components. This can be remedied by recomputing the quantities in two different ways:
**AND framework:** Here we consider a match only if *both* traces have SNR above the given threshold:

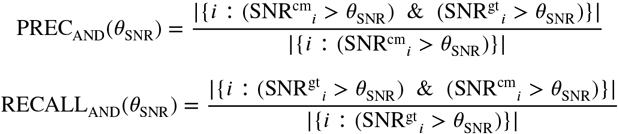
**OR framework:** Here we consider a match if *either* trace has SNR above the given threshold and its match has SNR above 0.

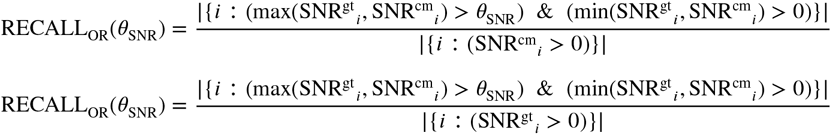 It is easy to show that

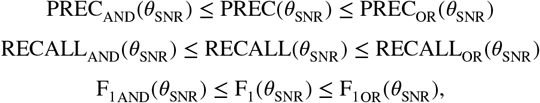

with equality holding for *θ*_SNR_ = 0. As demonstrated in Fig. 4d, these bounds are tight.

### Additional features of CaImAn

CaImAn contains a number of additional features that are not presented in the results section for reasons of brevity. These include:

#### Volumetric data processing

Apart from planar 2D data, CaImAn batch is also applicable to 3D volumetric data arising either from dense raster scanning methods, or from direct volume imaging methods such as light field microscopy (*Prevedel et al., 2014; Grosenick et al., 2017*).

#### Segmentation of structural indicator data

Structural indicators expressed in the nucleus and functional indicators expressed in the cytoplasm can facilitate source extraction and help identify silent or specific subpopulations of neurons (e.g., inhibitory). CaImAn provides a simple adaptive thresholding filtering method for segmenting summary images of the structural channel (e.g., mean image). The obtained results can be used for seeding source extraction from the functional channel in CaImAn batch or CaImAn online as already discussed.

#### Duplicate Detection

The ground truth obtained through the consensus process was screened for possible duplicate selections. To detect for duplicate components we define the degree of spatial overlap matrix *O* as

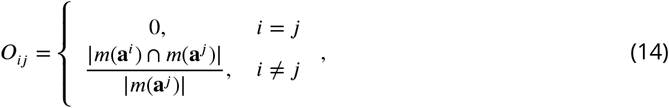

that defines the fraction of component *i* that overlap with component *j*, where *m*(·) is the thresholding function defined in (10). Any entry of *O* that is above a threshold *θ_o_* (e.g., *θ_o_* = 0.7 used here) indicates a pair of duplicate components. To decide which of the two components should be removed, we use predictions of the CaImAn batch CNN classifier, removing the component with the lowest score.

#### Extraction of Δ*F/F*

The fluorescence trace **f***_i_* of component *i* can be written as

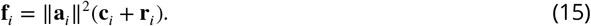

The fluorescence due to the component’s transients overlaps with a background fluorescence due to baseline fluorescence of the component and neuropil activity, that can be expressed as

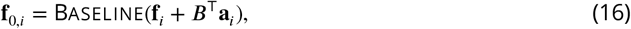

where Baseline: 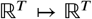 is a baseline extraction function, and ***B*** is the estimated background signal. Examples of the baseline extraction function are a percentile function (e.g., 10th percentile), or a for longer traces, a running percentile function, e.g., 10th percentile over a window of a hundred seconds^6^. To determine the optimal percentile level an empirical histogram of the trace (or parts of it in case of long traces) is computed using a diffusion kernel density estimator (*Botev et al., 2010*), and the mode of this density is used to define the baseline and its corresponding percentile level. The Δ*F/F* activity of component *i* can then be written as

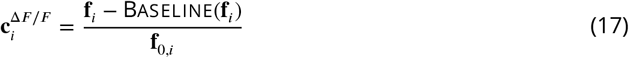

The approach we propose here is conceptually similar to practical approaches where the Δ*F/F* is computed by averaging over the spatial extent of an ROI (*Jia et al., 2011*) with some differences: i) instead of averaging with a binary mask we use the a weighed average with the shape of each component, ii) signal due to overlapping components is removed from the calculation of the background fluorescence, and iii) the traces have been extracted through the CNMF process prior to the Δ*F/F* extraction. Note that the same approach can also be performed to the trace ║**a***_i_*║^2^**c***_i_* that does not include the residual traces for each component. In practice it can be beneficial to extract Δ*F/F* traces prior to deconvolution, since the Δ*F/F* transformation can alleviate the effects of drifting baselines, e.g., due to bleaching. For the non-deconvolved traces **f***_i_* some temporal smoothing can also be applied to obtain more smooth Δ*F/F* traces.

## Acknowledgments

We thank B. Cohen, L. Myers, N. Roumelioti, and S. Villani for providing us with manual annotations. We thank V. Staneva and B. Deverett for contributing to the early stages of CaImAn, and L. Paninski for numerous useful discussions. We thank N. Carriero, I. Fisk, and D. Simon from the Flatiron Institute (Simons Foundation) for useful discussions and suggestions to optimize High Performance Computing code. We thank T. Kawashima and M. Ahrens for sharing the whole brain zebrafish dataset. Last but not least, we thank the active community of CaImAn users for their great help in terms of code/method contributions, bug reporting, code testing and suggestions that have led to the growth of CaImAn into a widely used open source package. A partial list of contributors (in the form of GitHub usernames) can be found in https://github.com/flatironinstitute/CaImAn/graphs/contributors (Python) and https://github.com/flatironinstitute/CaImAn-MATLAB/graphs/contributors (MATLAB^®^). The authors acknowledge support from following funding sources: A.G., E.A.P., J.F., P.G. (Simons Foundation). J.G., S.A.K., D.W.T. (NIH NRSA F32NS077840-01,5U01NS090541, 1U19NS104648 and Simons Foundation SCGB), P.Z. (NIH NIBIB R01EB022913, NSF NeuroNex DBI-1707398, Gatsby Foundation), J.T. (NIH R01-MH101198), F.N. (MURI, Simons Collaboration on the Global Brain and Pew Foundation).

## Supplemental Data

### Description of Supplemental Movies

**Movie 1:** Depiction of CaImAn online on a small patch of *in vivo* cortex data. Top left: Raw data. Bottom left: Footprints of identified components. Top right: Mean residual buffer and proposed regions for new components (in white squares). Enclosings of accepted regions are shown in magenta. Several regions are proposed multiple times before getting accepted. This is due to the strict behavior of the classifier to ensure a low number of false positives. Bottom right: Reconstructed activity.

**Movie 2:** Depiction of CaImAn online on a single plane of mesoscope data courtesy of E. Froudarakis, J. Reimers and A. Tolias (Baylor College of Medicine). Top left: Raw data. Top right: Inferred activity (without neuropil). Bottom left: Mean residual buffer and accepted regions for new components (magenta squares). Bottom right: Reconstructed activity.

**Movie 3:** Results of CaImAn online initialized by CaImAn batch on a whole brain zebrafish dataset. Each panel shows the active neurons in a given plane (top-to-bottom) without any background activity. See the text for more details.

## Algorithmic Details

In the following we present in pseudocode form several of routines introduced and used by CaImAn. Note that the pseudocode descriptions do not aim to present a complete picture and may refer to other work for some of the steps.

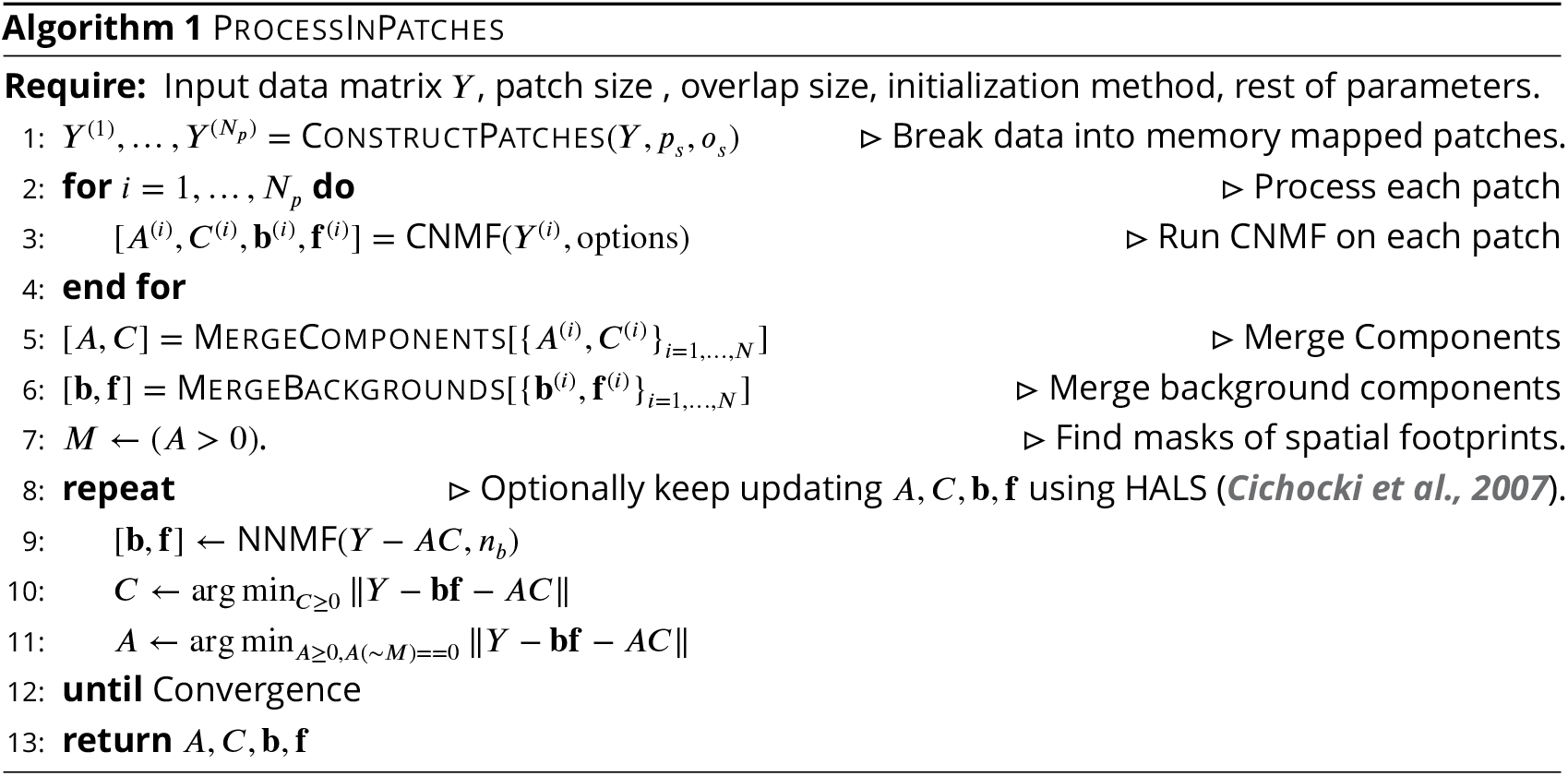

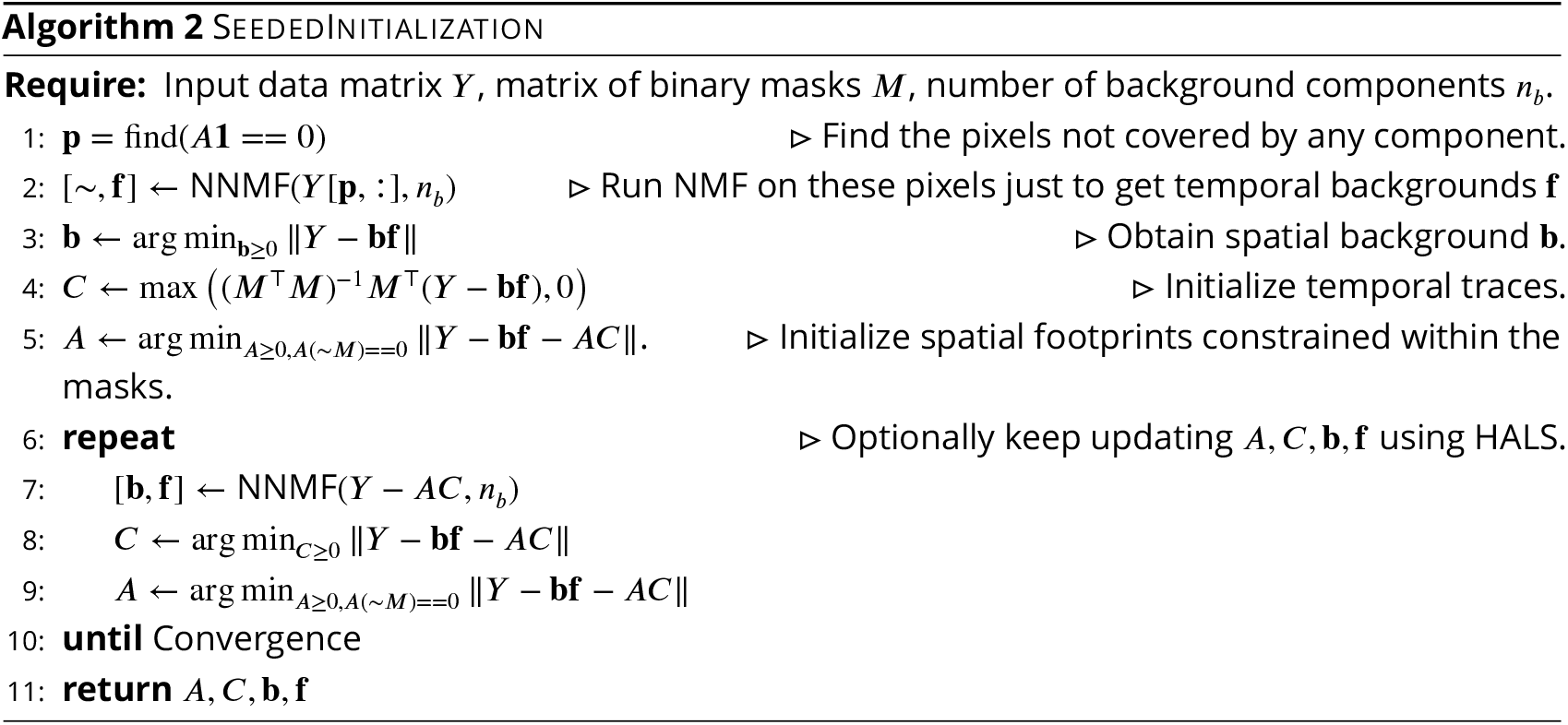

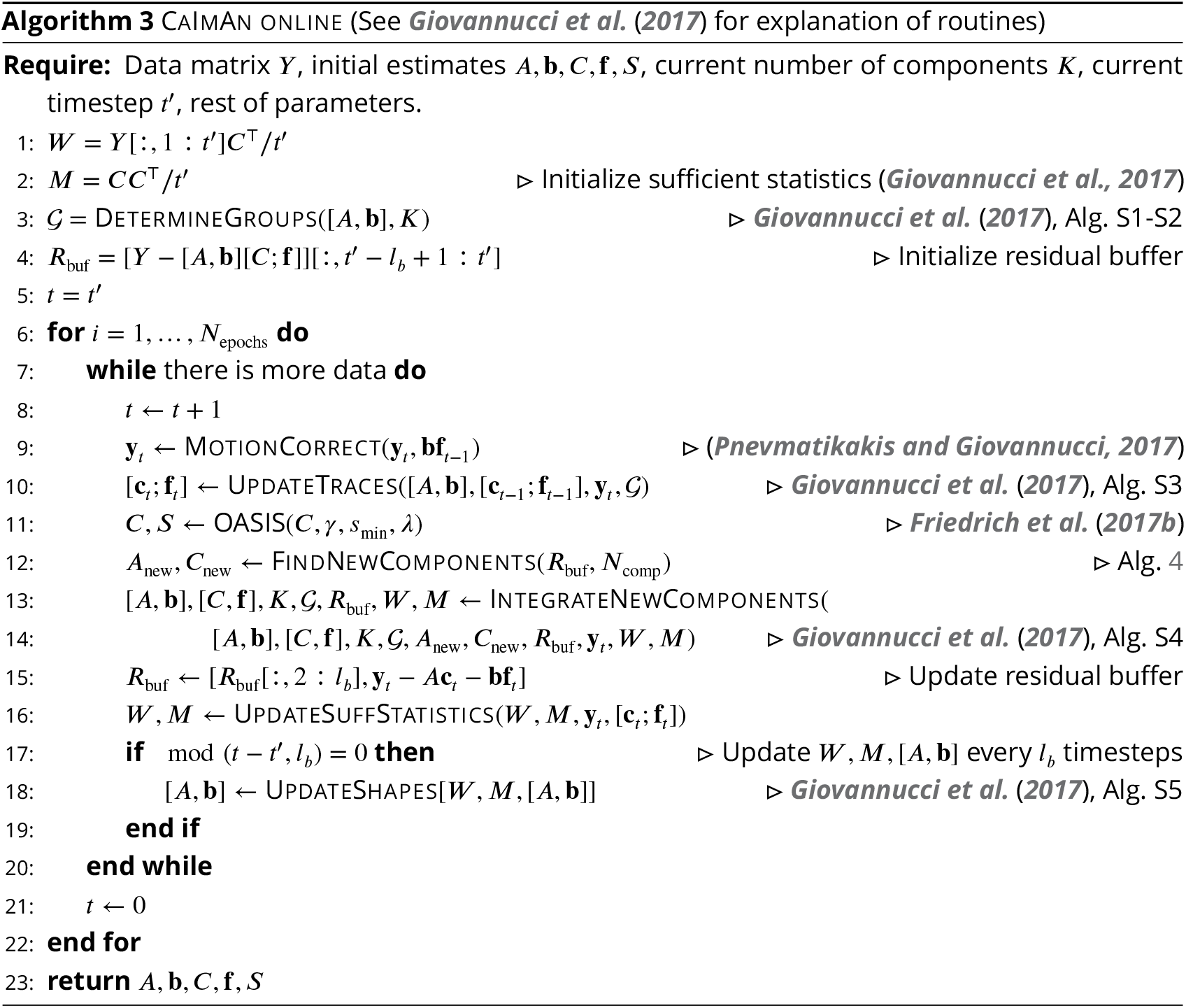

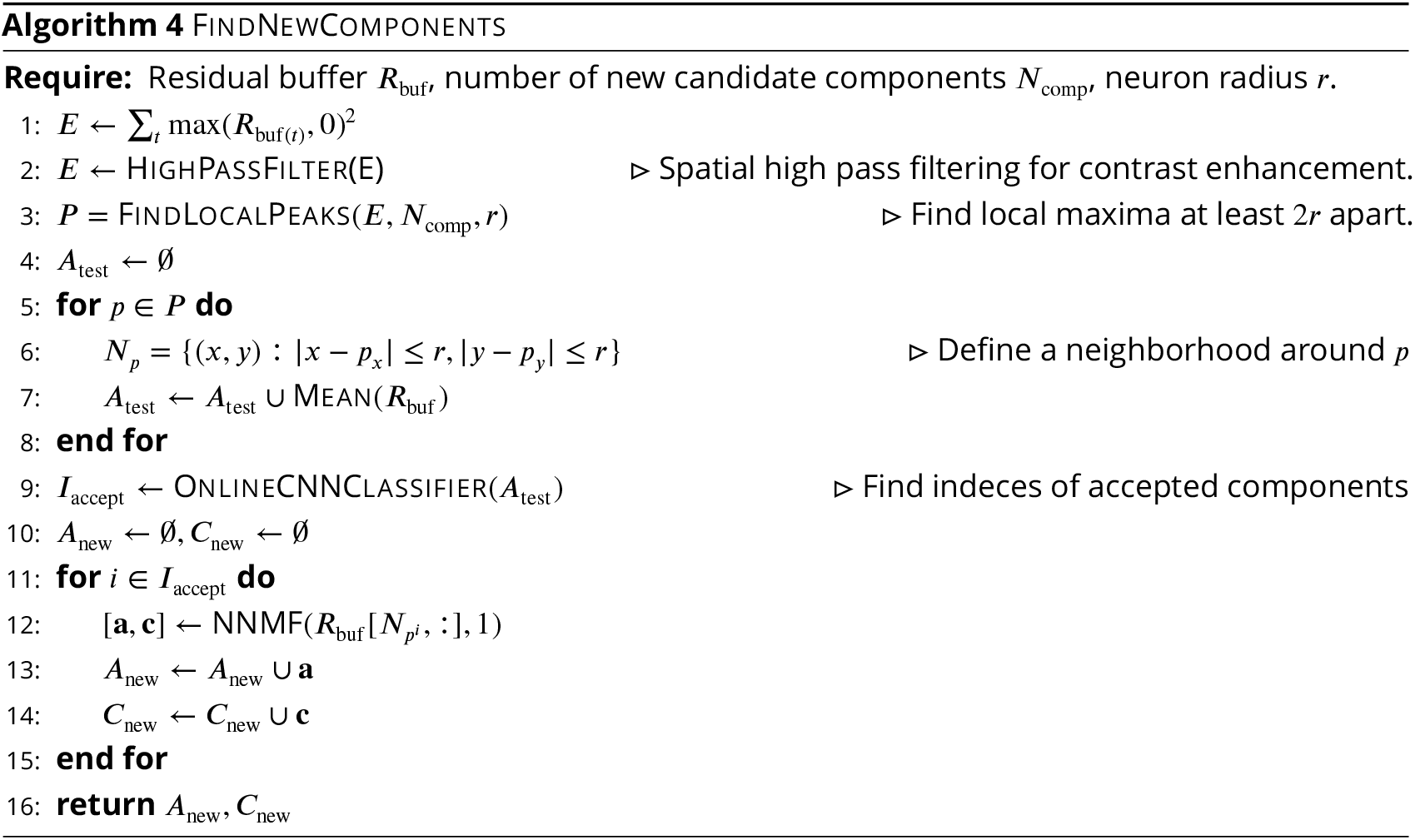

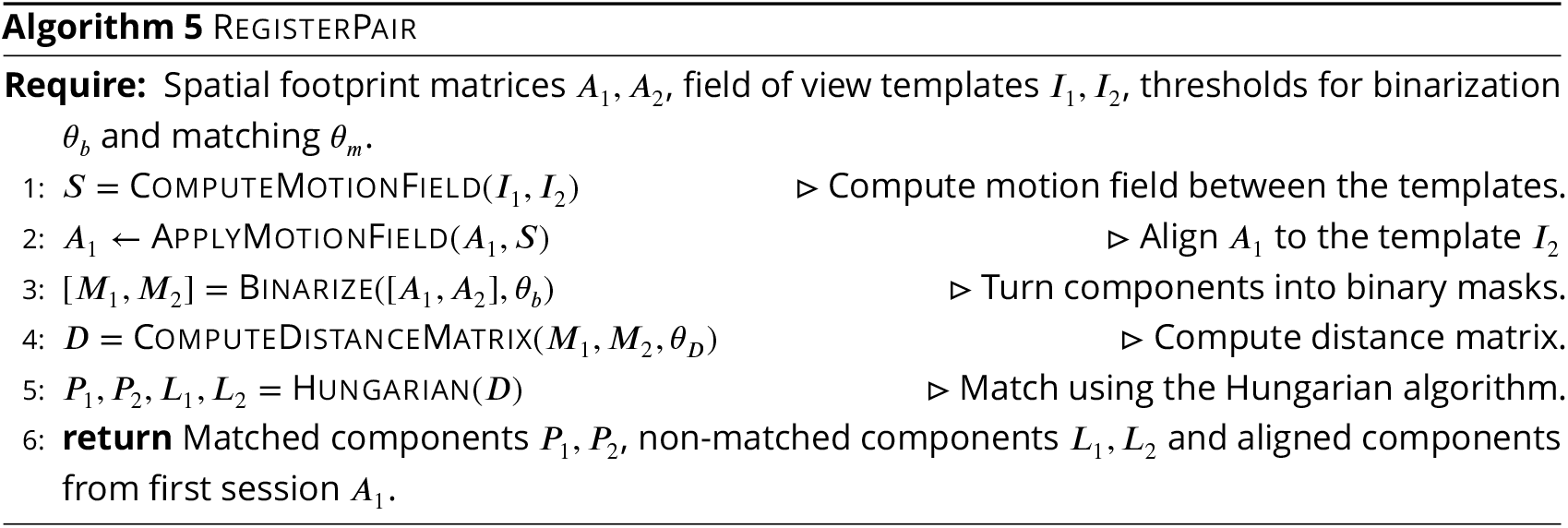

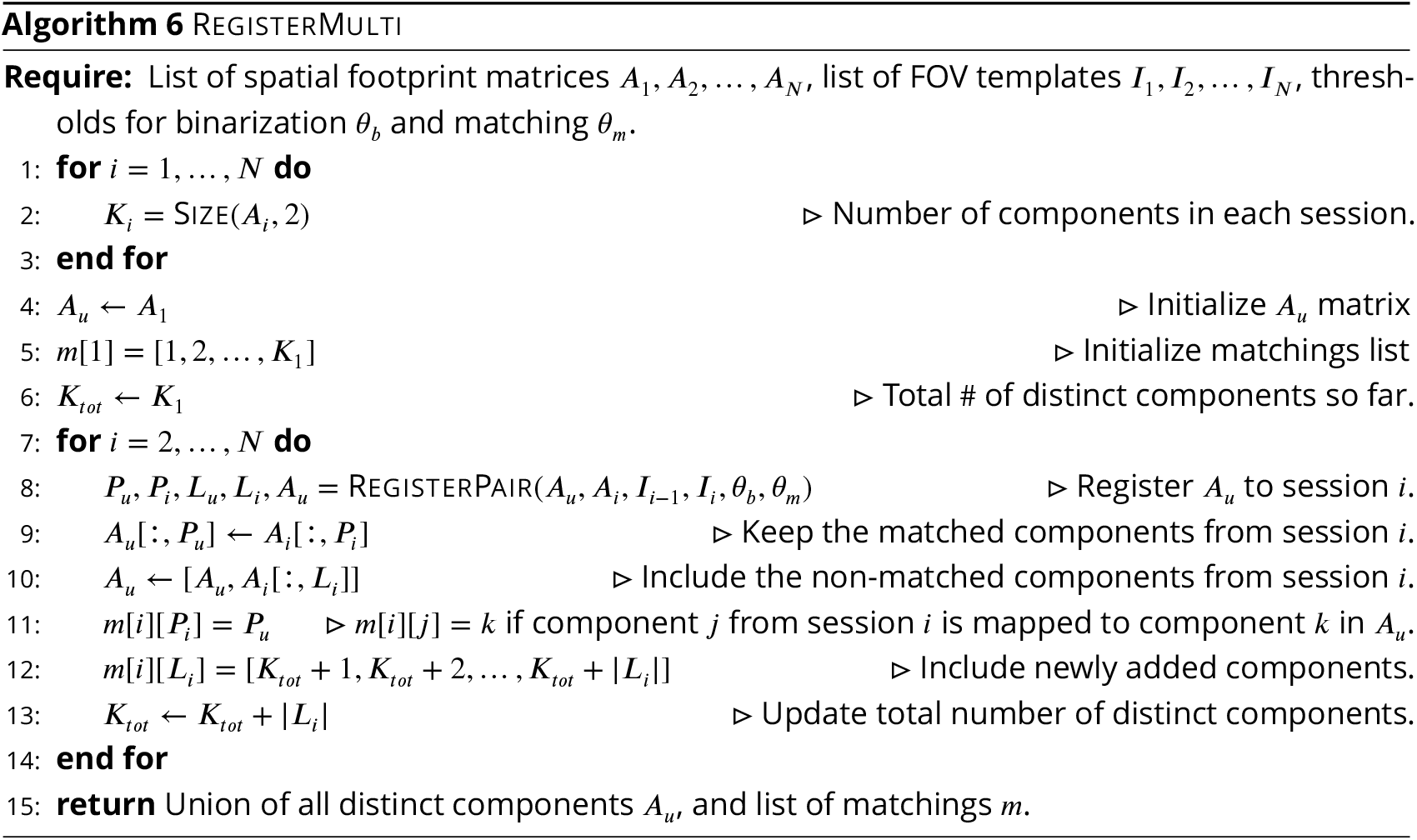

## Footnotes

1 Calculation performed on a 512×512 FOV imaged at 30Hz producing an unsigned 16-bit integer for each measurement.

2 Since proteins expressing the calcium indicator are confined outside the cell nuclei, neurons will appear as ring shapes, with a dark disk in the center.

3 It is possible that this process generated slightly biased results in favor of each individual annotators since the ground truth was always a subset of the union of the individual annotations.

4 All future development of CaImAn will be in Python 3, eventually rendering it incompatible with Python 2.x.

5 These precision and recall metrics are computed on different sets of neurons, and therefore strictly speaking one cannot combine them to form an *F*_1_ score. However, they can be bound from above by being evaluated on the set of matched and non-matched components where at least one trace is above the threshold (union of blue and pink zones in Fig. 4c) or below by considering only matched and non-matched components where both ground truth and inferred traces have SNR above the threshold (intersection of blue and pink zones in Fig. 4c). In practice these bounds were very tight for all but one dataset (Fig. 4d). More details can be found in *Methods and Materials (Performance quantification as a function of SNR)*.

6 Computing the exact running percentile function can be computationally intensive. To reduce the complexity we compute the running percentile with a stride of W, where *W* is equal or smaller to the length of the window, and then linearly interpolate the values.

